# Differences Among Fungal Microbiomes Associated with Tar Spot of Corn in Ecuador, Guatemala, and Indiana, USA

**DOI:** 10.1101/2025.02.07.637150

**Authors:** Wily R. Sic-Hernandez, Michael Gribskov, Alex G. Acosta, Andres Cruz, Jose L. Zambrano, Astrid J. Racancoj Coyoy, C. D. Cruz, Stephen B. Goodwin

## Abstract

Tar spot of corn, caused by the obligate fungal pathogen *Phyllachora maydis*, was previously found to be associated with *Monographella maydis* (syn. *Microdochium maydis*) and *Coniothyrium phyllachorae* in Mexico. While *M. maydis* was once believed to cause fisheye lesions and *C. phyllachorae* was described as a mycoparasite of *P. maydis*, their presence and roles in tar spot remain uncertain. This study hypothesizes that the tar spot-associated fungal microbiome varies geographically and includes fungal species with inconsistent associations with the disease. To test this, tar spot-infected leaves were collected in Guatemala, Ecuador, and the USA. This study shows that the distribution and abundance of these fungi exhibit a geographical pattern and are not consistently associated with tar spot lesions in all locations. We confirmed the presence of *Microdochium* in some parts of Ecuador and Guatemala but not in the USA. The role of *Microdochium* remains unclear and whether it can be a primary pathogen needs to be tested by additional research, as its occurrence and abundance depend more on geographical locations rather than the presence of fisheye lesions. *Coniothyrium* was detected at low abundance in samples from Ecuador and Guatemala but was absent in the USA. Additionally, we identified the likely mycoparasite *Paraphaeosphaeria* as highly prevalent across samples from all three countries. Microbiome diversity analysis grouped tar spot samples into four distinct clusters: two dominated by *P. maydis*, one primarily associated with *Microdochium*, and a fourth characterized by a combination of *P. maydis* and *Paraphaeosphaeria*.

## INTRODUCTION

Tar spot of corn is an emerging disease in the USA and Canada, with a broad geographic distribution in Central, South, and North America. This disease is caused by the obligate fungal pathogen *Phyllachora maydis* and is associated with *Coniothyrium phyllachorae* (Maublanc, 1904) and *Monographella maydis* (syn. *Microdochium maydis*) (Muller & Samuels, 1984) in Central and South America, but not in the USA. Earlier studies attributed fisheye lesions to *M. maydis*, although its pathogenic role remains debated, while *C. phyllachorae* was described as a mycoparasite of *P. maydis* (Hock et al., 1992).

In Mexico, Central, and South America, most reports of *M. maydis* and *C. phyllachorae* are based on morphological descriptions, with only a few confirmed by PCR. For instance, Hernandez (2018) isolated a fungus with *Microdochium*-like characteristics, and its ITS sequencing showed over 90% similarity to *Microdochium seminicola* and *Monographella* sp., along with sequences showing more than 93% similarity to *Coniothyrium*. Herrera et al. (2017) also reported *Curvularia lunata* in tar spot lesions in Mexico via PCR. A study conducted in Ecuador, however, isolated neither *C. phyllachorae* nor *M. maydis* from tar spot-infected leaves (Roman et al., 2018). Furthermore, analysis of fungi isolated from fisheye lesions collected in Mexico did not identify *Monographella* or *Microdochium* but instead classified the cultured fungi within the *Fusarium equiseti-incarnatum* species complex (J. M. Luis et al., 2023; McCoy et al., 2019). Sequence-based microbiome analyses in the USA similarly showed low abundance or absence of *C. phyllachorae*, but instead detected high levels of *Paraphaeosphaeria* (Caldwell et al., 2023; McCoy et al., 2019), another fungus that can be a mycoparasite. Therefore, the presence of *C. phyllachorae* and *M. maydis* in tar spot lesions remains uncertain, and they may not be present in all locations where *P. maydis* is responsible for causing the disease.

The diverse environments where tar spot has been reported also may influence the presence of *C. phyllachorae* and *M. maydis*. Tar spot has a broad distribution, being reported in 19 countries across the Americas (Valle-Torres et al., 2020) and 18 states within the USA (Crop Protection Network, 2024). Optimal conditions for natural infections are hot, humid environments (Liu, 1973) but tar spot has become an emergent disease in non-tropical corn regions with temperate or even cold weather (Racancoj-Coyoy et al., 2024) even though it likely coevolved with corn for thousands of years in the tropical zones of Latin America (Mottaleb et. al. 2018). In controlled environments, symptoms develop at temperatures between 18-24°C with relative humidity between 80-100% for initial infection and 20-100% for disease progression (Breunig et al., 2023; Solórzano, et al., 2023; Jimenez-Beitia et al., 2025).

Despite this, tar spot is found across diverse regions. For instance, Ecuador has three regions characterized by distinct ecological conditions: coastal lowlands, the Amazon rainforest, and the highlands. The coastal lowland is a relatively flat region at elevations of less than 200 feet above mean sea level (FAMSL), bordered by huge alluvial fans in the east, covered by a tropical rain forest in the north, and transitioning to an arid region in the south with annual average temperatures of 26 °C (78.8 °F). The highland region is composed of the Cordillera Occidental and the Cordillera Central divided by ten main basins formed by intense volcanic activity, with an elevation between 8,200 to 9,500 FAMSL and an annual average temperature of 12 °C (53.6 °F). The Amazon Rainforest is covered by dense tropical rainforest with temperatures between 13 to 29 °C (55.4 to 84.2 °F) (Brawer, 1992; Millan, et al. 2008; Morán-Tejeda, et at. 2016) and elevations from 800 to 1100 FAMSL. Notably, tar spot is present in all these regions, highlighting its ability to thrive under varying environmental conditions.

Guatemala is a country with high biodiversity, with average temperatures ranging from 9.5 to 22.6 °C (49 to 72 °F) in the Central Highlands and up to 14.3 to 29 °C (57.7 to 84.2 °F) in areas with more tropical conditions. Elevations range from 3100 to 7500 FAMSL. This makes it an important location for studying the tar spot-associated microbiome, in comparison with Ecuador. Furthermore, the appearance of fisheye lesions and stromata differs between geographical locations. For instance, in the USA, tar spot manifests mostly as rounded or oblong black, shiny stromata, occasionally surrounded by necrotic halos (Solórzano, Cruz, et al., 2023). In Mexico, lesions are described as round, glossy black stromata with narrow chlorotic borders, which eventually become necrotic (Bajet et al., 1994). In Guatemala, lesions are characterized by shiny black spots with an oval to semi-oval shape, ranging from 0.5-2 mm in diameter and up to 10 mm in length (López et al., 2011).

The diverse environmental conditions where tar spot is present could impact the microbiome associated with the disease and influence the appearance of tar spot lesions. These environmental variations also could explain why *Monographella maydis* has not been identified in the USA despite being commonly reported in other countries. The presence of *Monographella maydis*, *Coniothyrium phyllachorae*, and other fungi such as *Paraphaeosphaeria* in tar spot lesions may be location dependent. To test the hypothesis that fungal microbiomes associated with tar spot vary by location, we used Next-Generation Sequencing (NGS) to identify fungal communities associated with tar spot in Guatemala, Ecuador, and the USA. NGS allows for the identification of organisms based on their DNA sequences, eliminating the need to rely solely on morphological descriptions and reducing the risk of misidentification. This method has been used to study fungal communities in tar spot lesions in Michigan and Indiana (Caldwell et al., 2023; McCoy et al., 2019) but has yet to be applied to other regions.

By identifying the fungal microbiome associated with tar spot lesions across these different regions, we aim to understand better the fungi required to induce tar spot and fisheye lesions. Additionally, this research could reveal microbiota that interact with *Phyllachora maydis* in specific locations with potential for biological control of the disease. This study focuses on identifying *Phyllachora maydis*, *Coniothyrium*, *Monographella*, and *Paraphaeosphaeria*, as they have been reported to play a role in tar spot lesions, and aims to unravel the geographical variation in fungal microbiomes associated with tar spot and its implications for disease ecology and management.

## MATERIALS AND METHODS

### Sample collection

Tar spot-diseased corn leaves were collected in Ecuador (2021 and 2022), Guatemala (2022), and the USA (2021 and 2023). The selection of sampling sites was based on disease prevalence and agroecological diversity. In Ecuador, three fields were sampled in the Coastal lowlands, two in the Amazon, and two in the Highlands region, while in Guatemala seven fields in the Highlands region were sampled (Table 1). In both countries, infected leaves were collected from different corn varieties and physiological stages but all at mid-canopy. At least three infected leaves were collected in each field and transported in individual paper bags. Leaves were dried in an artisan press using newspaper and cardboard and stored at room temperature until leaves were completely dry (indicated by breaking cleanly when bent, typically within 6 to 11 days). A portion of 6 inches was cut from each leaf and placed in a paper envelope with silica gel packets. Envelopes were placed in Ziploc bags and shipped to the USA under USDA permit number 526-23-102-80391. Upon arrival, samples were stored at 4 degrees Celsius until DNA extraction. In the USA, samples were collected at the Pinney Purdue Agricultural Center (PPAC) located in LaPorte, Indiana, during the summers of 2021 and 2023. A total of ten leaves were collected in 2021 and 12 in 2023. Samples collected in 2021 were dried and stored in an artisan press until DNA extraction. Samples collected in 2023 were folded in aluminum foil, transported with packs of dry ice, and stored at −20 degree Celsius until DNA extraction.

**Table 1.** Summary information about the locations where tar spot samples were collected.

| Country | Region | State | Location | Corn Type | Date of collection | Storage method | Elevation (FAMSL)* | Latitude, longitude decimal degrees | Total samples |
| --- | --- | --- | --- | --- | --- | --- | --- | --- | --- |
| USA | Midwest | Indiana | La Porte | Hybrid | Aug-2021 | Dry | 760 | 41.453623, -86.941487 | 10 |
| USA | Midwest | Indiana | La Porte | Hybrid | Sept-2023 | Frozen | 760 | 41.453623, -86.941487 | 12 |
| Guatemala | Central Highlands | Chimaltenango | Técpan | Landrace | Oct-2022 | Dry | 7,496 | NA | 2 |
| Guatemala | Central Highlands | Chimaltenango | Parramos | Landrace | Oct-2022 | Dry | 6,417 | 14.583367, -90.837924 | 1 |
| Guatemala | Central Highlands | Chimaltenango | ICTA | Variety | Oct-2022 | Dry | 5,905 | 14.631865, -90.804703 | 3 |
| Guatemala | Central highlands | Sacatepéquez | Antigua Guatemala | NA<sup>+</sup> | Oct-2022 | Dry | 5,019 | 14.666144, -90.70914 | 3 |
| Guatemala | Central Highlands | Baja Verapaz | Rabinal | NA | Aug-2022 | Dry | 3,192 | 15.119361, -90.531056 | 3 |
| Guatemala | Central Highlands | Baja Verapaz | Cubulco | NA | Sept-2022 | Dry | 3,264 | NA | 3 |
| Guatemala | Central highlands | Chimaltenango | Patzún | Hybrid | Oct-2022 | Dry | 7,316 | 14.66088, -90.998 | 3 |
| Ecuador | Amazon | Sucumbios | El Eno | Hybrid | Nov-2022 | Dry | 1033 | -0.035, -76.644 | 5 |
| Ecuador | Amazon | Orellana | La Joya Sachas | Hybrid | Nov-2022 | Dry | 820 | -0.441, -76.568 | 5 |
| Ecuador | Coastal Lowlands | Los Rios | Mocache | Hybrid | Aug-2022 | Dry | 150 | -1.185774, -79.535656 | 3 |
| Ecuador | Coastal Lowlands | Los Rios | Vince | Hybrid | Aug-2022 | Dry | 55 | -1.365182, -79.554782 | 4 |
| Ecuador | Coastal Lowlands | Los Rios | Pueblo Viejo | Hybrid | Dec-2021 | Dry | 26 | -1.560072, -79.501446 | 3 |
| Ecuador | Andean Highlands | Pichincha | Quito | Landrace | Aug-2022 | Dry | 8,530 | -0.140117, -78.436584 | 9 |
| Ecuador | Andean Highlands | Imbabura | Ibarra | Landrace | Aug-2022 | Dry | 7,749 | 0.356077, -78.206986 | 4 |

\* Feet above mean sea level.
<sup>+</sup> Data not available.

### DNA extraction and sequencing

From each leaf or leaf portion, six discs (18 mm in diameter) were excised using a sterilized Cork borer, with each disc containing one stroma. However, in some samples from Indiana, USA, the stromata were small and clustered together, causing the discs to contain more than one stroma. The leaf discs were vortexed at high speed in sterile water for 10 seconds to remove debris, then dried with paper towels. DNA extraction was performed using the E.Z.N.A.® Plant DNA DS Kit (D2411, Omega Bio-tek, Inc. Norcross, GA), following the manufacturer’s instructions with minor modifications. The six leaf discs from a single leaf were ground in a mortar and pestle with liquid nitrogen. The resulting powder was transferred to a Synergy Homogenization Tube (SKU SYNP 100-02, OPS Diagnostics, Lebanon, NJ) with 750 μl of CSPL buffer from the E.Z.N.A. kit, and the tissue was homogenized using a bead beater at high speed for 5 minutes. DNA quality was assessed using a spectrophotometer and a Qubit Fluorometer, ensuring an OD260/280 ratio between 1.8 to 2.0 and a concentration greater than 200 ng.

Library construction, quality control, and sequencing were conducted by Novogene (Sacramento, CA). The Internal Transcribed Spacer regions I and II (ITS1 and ITS2) were amplified using PCR. For ITS1, the forward primer GGAAGTAAAAGTCGTAACAAGG and the reverse primer GCTGCGTTCTTCATCGATGC were used. For ITS2, the forward primer GCATCGATGAAGAACGCAGC and the reverse primer TCCTCCGCTTATTGATATGC were employed. Barcodes were added to all primers to identify sequences from individual samples. PCR products of the appropriate size were selected using 2% agarose gel electrophoresis. Equal amounts of PCR products from each sample were pooled, end-repaired, A-tailed, and ligated with Illumina adapters. Library quality was assessed and quantified using qPCR, and quantified libraries were pooled based on effective library concentrations and the required data output. Sequencing was performed on a paired-end Illumina platform, generating 250-b paired-end raw reads. Paired-end reads were assigned to their corresponding samples based on their unique barcodes, and the barcodes and primer sequences were removed computationally.

### Taxonomic annotation

Raw sequence reads were analyzed using Quantitative Insight Into Microbial Ecology (QIIME2) v.2024.2 (Bolyen et al., 2019). During the DADA2 v1.26.0 denoising step (Callahan et al., 2016), ITS1 forward and reverse sequences were trimmed at position 20 bp and truncated at position 200 bp and 110 bp, respectively, to ensure that the sequences had a quality greater than Q20. Similarly, ITS2 forward and reverse sequences were trimmed at position 20 bp and truncated at position 214 bp and 175 bp, respectively, to achieve the same quality threshold (Q > 20). Sequences were then filtered, dereplicated, denoised, and the forward and reverse sequences were merged, followed by the removal of chimeras. The percentage of reads removed was 42.6% and 19.1% from a total of 14,916,896 and 14,121,512 sequencing reads for ITS1 and ITS2, respectively. DADA2 subsequently was used to cluster the sequences into amplicon sequence variants (ASVs) with a single-nucleotide resolution, resulting in 2,885 and 3,921 ASVs for ITS1 and ITS2, respectively. The taxonomy of each ASV was assigned using the UNITED database version 10.0 released on April 4^th^, 2024 (https://dx.doi.org/10.15156/BIO/2959337). Sequences annotated as mitochondrial and chloroplast were removed. The number of reads per sample ranged from 20,117 to 260,131 for ITS1 and 41, 418 to 192, 791 for ITS2. ITS1 sequences did not provide enough taxonomic resolution to identify *P. maydis* (Supplementary Fig. 1), therefore only ITS2 sequences were considered for further analysis. At the lowest number of sequencing reads per sample, rarefaction curves for the other samples began to reach an asymptotic plateau (Supplementary Fig. 2), indicating sufficient sequencing depth. Consequently, no samples were excluded from the analysis.

QIIME2 outputs were imported to R using the package “phyloseq” version 1.48.0. (McMurdie & Holmes, 2013) to create bar plots using the package “ggplot2” version 3.5.1 (Wickham, 2016). For bar plotting, if an ASV was not classified at the species level, it was retained and assigned to the lowest taxonomic level for which classification was available (e.g., genus) to ensure it was not excluded from the analysis. As a result, the species bar plots show taxa labels such as “uncl. Pleosporales” which stands for unclassified Pleosporales. Finally, data were filtered to show only the 30 most abundant taxa; as a result, bars do not reach 100% relative abundance.

### Population diversity analysis

Alpha and beta diversity metrics were estimated in QIIME2 using the “diversity” plugin. The alpha diversity of fungal communities in tar spot of corn was measured based on the number of species (richness) in a sample and their abundance distribution (evenness). Richness was measured with Faith’s phylogenetic index (Faith, 1992), which considers the number of species and their evolutionary relationships. Evenness was evaluated with Pielou’s evenness index (Pielou, 1966), which measures how the species are distributed in the community. We also used the Shannon index (Shannon, 1948) which accounts for both richness and evenness, capturing tar spot microbiome composition comprehensively. Alpha diversity indexes were used to compare diversity across countries, geographical regions, and locations. Beta diversity was calculated using the Bray-Curtis dissimilarity index (Bray & Curtis, 1957) to compare fungal abundance across samples and the Weighted Unifrac index (Lozupone et al., 2007) to measure differences in fungal communities based on phylogenetic relationships and species abundance. Beta-diversity indices were applied to assess differences between countries and geographical regions.

QIIME2 outputs were imported into R to create box plots for alpha indexes and principal coordinate analysis (PCoA) plots for beta-diversity indexes. Statistical analysis of the alpha indexes was performed in R using the package “stats” version 4.4.0. First, the Kruskal-Wallis rank sum test was used to assess significant differences between groups, followed by pairwise comparisons using the Wilcoxon rank sum exact test. Statistical analysis of the beta indexes was performed in R using the function “adonis2” from the package vegan version 2.6.6.1 to perform a permutational multivariate analysis of variance test (PERMANOVA).

### Differential abundance analysis between fisheye and no fisheye samples

The Analysis of Compositions of Microbiomes with Bias Correction (ANCOM-BC) algorithm (Lin & Peddada, 2020) was used to identify species that are differentially abundant between fisheye and no fisheye samples. The analysis was conducted in QIIME2 using the plugin “q2-composition” version 2024.2.0. First, species with a relative abundance of less than 0.01% were discarded; this removed 1.98% of the total reads. A second filter was applied to exclude those species present in less than 20% of the total samples. A bar plot was created to show species that were differentially abundant with p values less than or equal to 0.05 indicating significant differences.

### Fungal correlations

Correlations were assessed among the 20 most abundant fungal species using Spearman’s rank correlation coefficients, calculated with the cor() function in R. The statistical significance of the correlation matrix was tested using the cor.mtest() function from the corrplot package at a 95% confidence level. The results were visualized in a correlation matrix plot.

The ASVs with the highest sequence read counts of the 20 most abundant species in tar spot were compared to sequences in the UNITED database and NCBI’s core nucleotide database to determine the percentage of identity and nucleotide differences. For example, 4,004,034 sequences were classified as *P. maydis* and grouped into 140 ASVs; however, three ASVs accounted for 95.3% of the total sequences assigned to this taxon so only these three were considered in further analyses. The first of the *P. maydis* ASVs matche Group 3 of Broders et al. (2022) with an E-value of 9e^−141^ and a 100% identity. Group 3 is comprised of 11 *Phyllachora* species, including *P. maydis*. Samples in this group were collected from different countries and years from 1917 to 2019 (Broders et al., 2022). This ASV also matched the other groups but with a lower E-value and an identity between 88% to 96%. The second and third *P. maydis* ASVs matched with Group 2 of Broders et al., (2022) with an E-value of 9e^−139^ and anidentity of 99% (1 bp of difference). Group 2 is comprised of only *P. maydis* samples collected in the USA, Mexico, Puerto Rico, and Colombia from 1917 to 2019 (Broders et al., 2022). These ASVs also matched the other groups with lower E-values and an identity between 85% to 99% (2 bp of difference). Since some ASVs could not be classified to species level, they are retained at the lowest taxonomic level to which classification was possible. As a result, the results include taxa at the genus and family levels.

## RESULTS

### Fungal microbiomes associated with tar spot

*Phyllachora maydis*, *Monographella* (syn. *Microdochium*) *maydis* and *Coniothyrium phyllachorae* have been reported from tar spot lesions previously. The results of this study confirm the presence of all three species in tar spot lesions. Additionally, we observed a high relative abundance of *Paraphaeosphaeria* sp. across all three countries. However, species presence and abundance varied significantly between countries, geographical regions, and even locations. We found that an ASV with its highest match to *Microdochium seminicola* was highly present in Ecuador and Guatemala but was almost absent in the USA, while *Coniothyrium* was present only in Ecuador and Guatemala (Table 2).

**Fig. 1.**
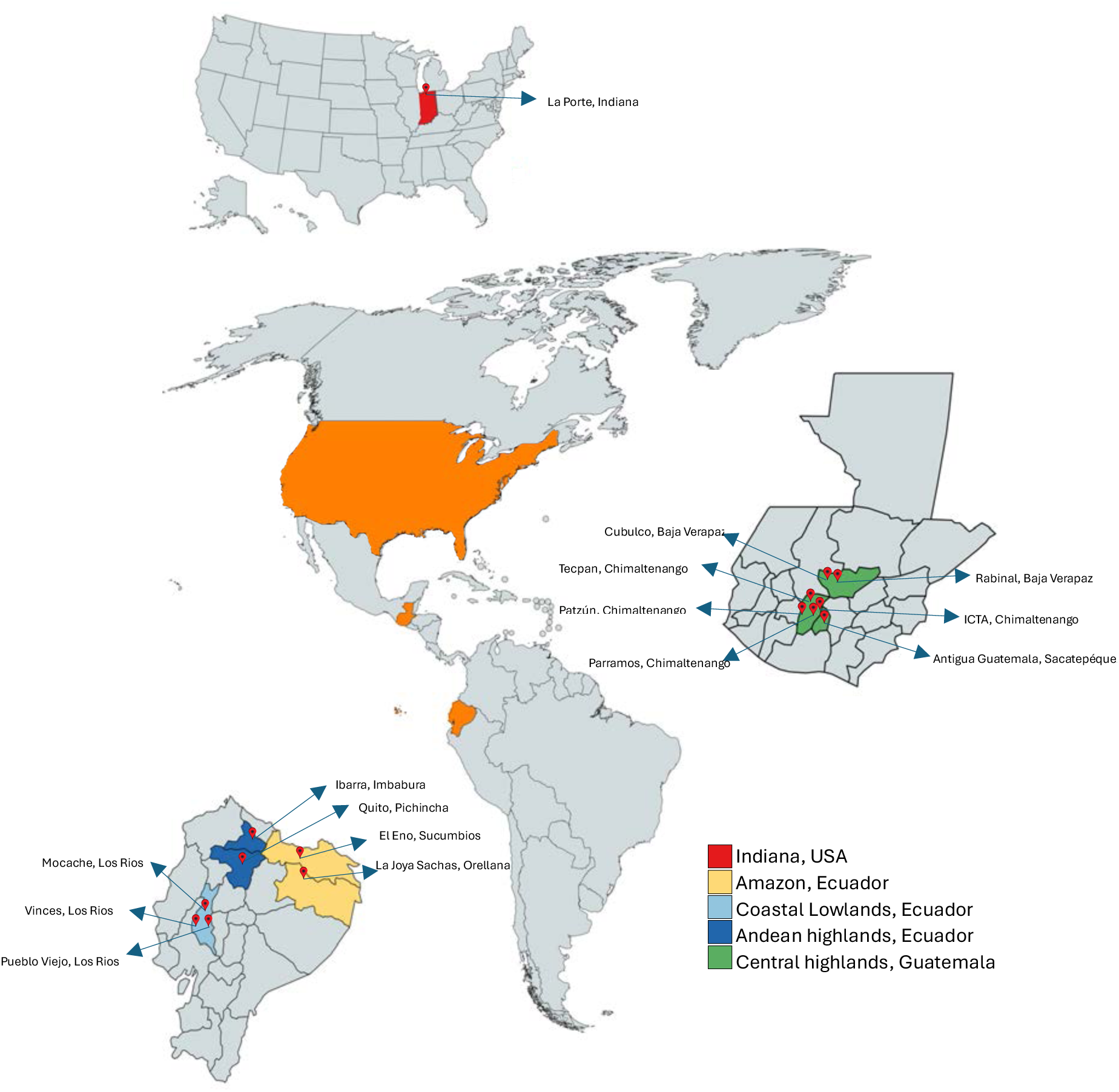
Sample locations in the USA, Ecuador, and Guatemala. In Ecuador, states marked in yellow are located in the Amazon, those in light blue are in the Coastal Lowlands, and those in dark blue are in the Andean Highlands. In Guatemala, states colored in dark green are located in the Central Highlands. In the USA, the state colored in red is located in the Midwest.

**Table 2.** Relative abundance of fungal species present in tar spot lesions in Ecuador, Guatemala, and the USA locations.

| Country | Region | Location | *Phyllachora*<br>*maydis* (%) | *Paraphaeosphaeria*<br>*sp.* (%) | *Microdochium*<br>*seminicola* (%) | *Coniothyrium*<br>*sidae* |
| --- | --- | --- | --- | --- | --- | --- |
| USA | Indiana | La Porte | 64.3 | 7.0 | <0.0 | 0.0 |
| Ecuador | Amazon | El Eno | 9.6 | <0.0 | 28.5 | 0.0 |
| Ecuador | Amazon | La Joya Sacha | <0.0 | <0.0 | 37.7 | 0.0 |
| Ecuador | Coastal Lowlands | Mocache | 1.2 | 0.1 | 68.1 | <0.0 |
| Ecuador | Coastal Lowlands | Vinces | 5.0 | 0.1 | 46.3 | <0.0 |
| Ecuador | Coastal Lowlands | Pueblo Viejo | 2.5 | 42.5 | 0.1 | 0.0 |
| Ecuador | Andean Highlands | Ibarra | 26.0 | 59.8 | 0.1 | <0.0 |
| Ecuador | Andean Highlands | Quito | 50.9 | 38.9 | 0.4 | <0.0 |
| Guatemala | Central Highlands | Técpan | 71.9 | <0.0 | 0.6 | <0.0 |
| Guatemala | Central Highlands | Chimaltenango | 26.9 | 1.3 | 0.9 | <0.0 |
| Guatemala | Central Highlands | Parramos | 27.1 | 63.6 | 3.4 | 0.0 |
| Guatemala | Central Highlands | Patzún | 17.9 | 38.4 | 0.4 | 0.0 |
| Guatemala | Central Highlands | Antigua<br>Guatemala | 68.2 | 0.9 | 0.2 | 0.0 |
| Guatemala | Central Highlands | Cubulco | 3.9 | 0.1 | 17.1 | <0.0 |
| Guatemala | Central Highlands | Rabinal | 0.2 | <0.0 | 31.8 | <0.0 |

In samples from Indiana, *Phyllachora maydis* was the most abundant species (64.3%), followed by *Paraphaeosphaeria* sp. (7.0%), *Alternaria alternata* (5.9%), and *Nigrospora* sp. (4.7%). Only six sequences of *Microdochium* spp. were found in two out of 22 samples collected in this country, representing a mere relative abundance of 0.0002%. The six sequences are clustered into two different ASVs, with one present in the three countries and the other only in the USA. Meanwhile, *Coniothyrium* was absent from all samples from Indiana.

In Ecuador four taxa predominated in tar spot lesions: *Microdochium seminicola* (22.7%), *Paraphaeosphaeria* sp. (21.4%), *Phyllachora maydis* (18.6%), and *Russulales* sp. (8.3%) (Fig. 2). In this case, *M. seminicola* was the closest database hit. However, it varied from zero to 16 nucleotide positions (94 to 100% identity) when compared with different accessions of that species, so it likely represents a different, unnamed species or at least one that is not present in the database. For simplicity it will be referred to as *M. seminicola* here, but its correct identity will need to be determined by additional analysis in the future. The relative abundance of *M. seminicola* exceeded that of *Phyllachora maydis* in the Amazon and Coastal Lowlands regions of Ecuador, but was almost absent in the Andean Highlands, where *Phyllachora maydis* and *Paraphaeosphaeria* sp. were the dominant species (Fig. 3).

**Fig 2.**
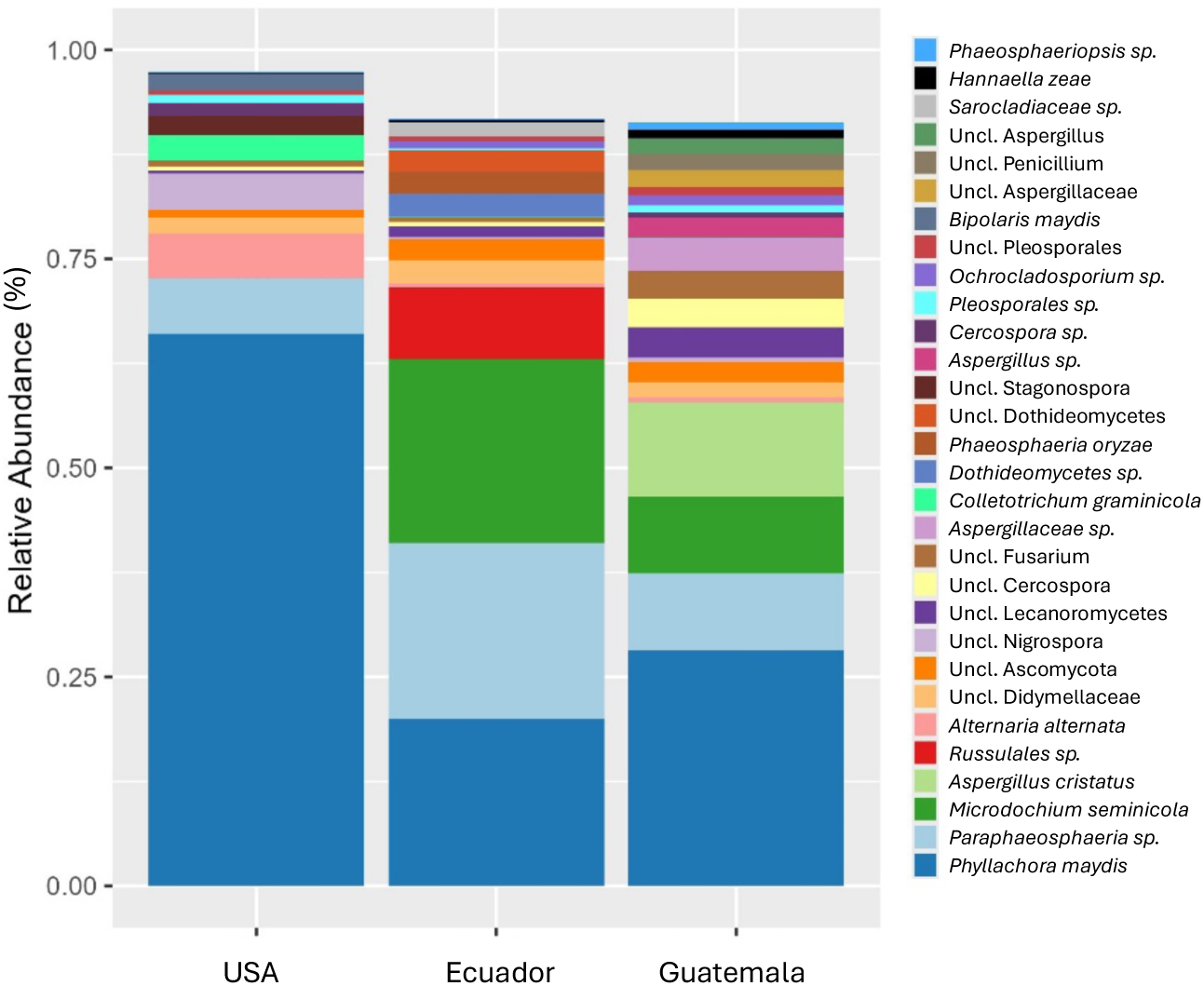
The 30 most abundant fungal species in tar spot lesions collected in the USA, Ecuador, and Guatemala. Uncl.: Unclassified species that belong to a taxonomic group but could not be assigned to a specific species. Bars do not reach 100% as species beyond the 30 most abundant are not displayed.

**Fig 3.**
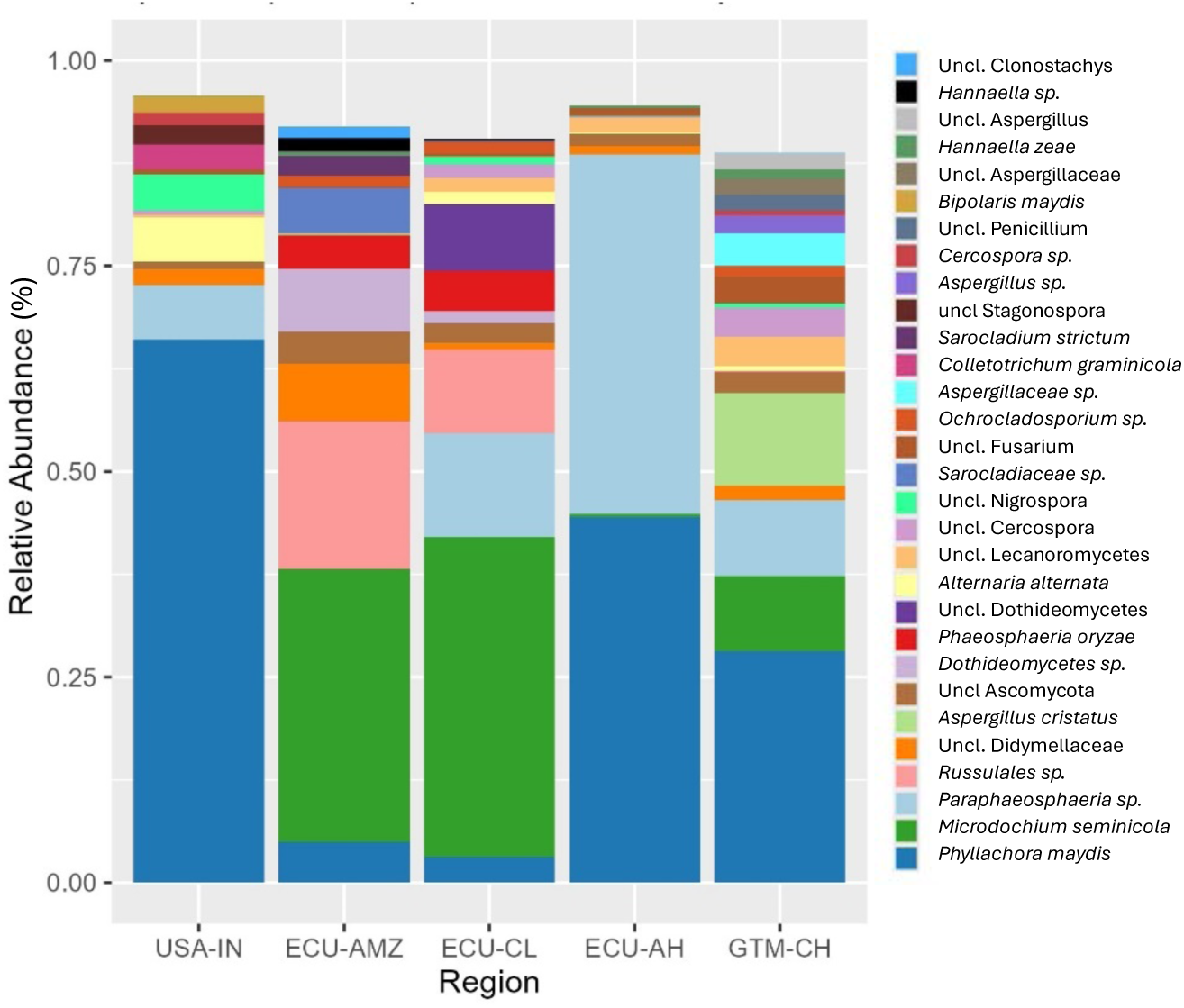
The 30 most abundant fungal species in tar spot lesions by region. USA-IN: Indiana, ECU-AMZ: Ecuadorian Amazon, ECU-CL: Costal Lowlands of Ecuador, ECU-AH: Andean Highlands of Ecuador, and GTM-CH: Central Highlands of Guatemala. Uncl.: Unclassified species that belong to a taxonomic group but could not be assigned to a specific species. Bars do not reach 100% as species beyond the 30 most abundant are not displayed.

In the Ecuadorian Amazon region, *Microdochium seminicola* was the most abundant taxon, while *Phyllachora maydis* was scarce. In fact, *P. maydis* was found in only one sample from La Joya Sachas (Fig. 4). *Paraphaeosphaeria sp.* in this region is also low (0.01%) and found in only six out of 10 samples. Interestingly, *Russulales* sp. (35.9%) was present in a high abundance in the location El Eno, surpassing *Microdochium seminicola* (28.5%) and *Phyllachora maydis* (9.7%). Other taxa, including unclassified species from *Didymellaceae*, *Dothideomycetes* and *Sarocladiaceae*, appeared in moderate abundances in this region.

**Fig 4.**
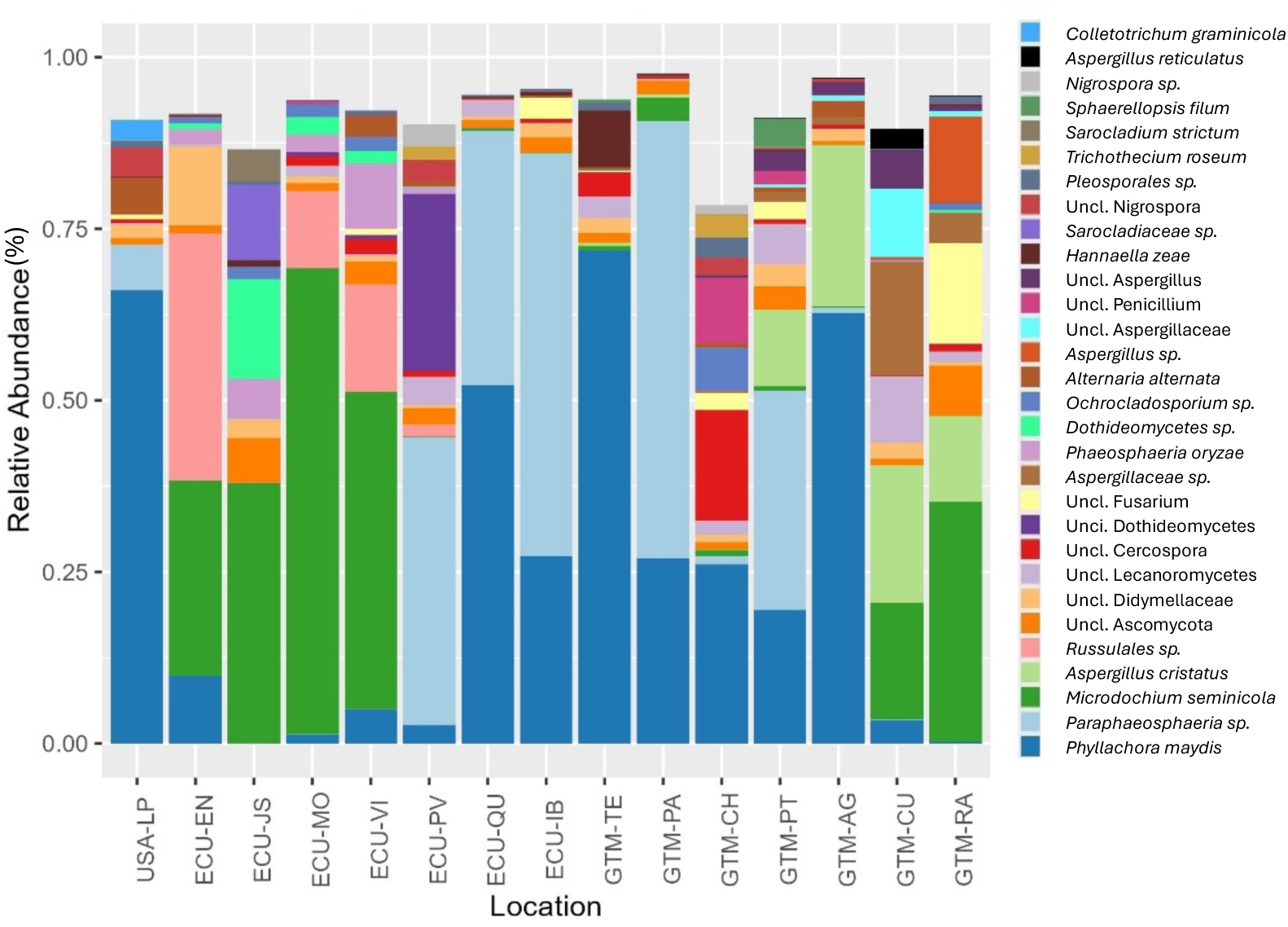
The 30 most abundant fungal species in tar spot lesions by location. USA-LP: La Porte, Indiana; ECU-EN: El Eno, Ecuadorian Amazon; ECU-JS: La Joya Sachas, Ecuadorian Amazon; ECU-MO: Mocache, Coastal Lowlands of Ecuador; ECU-VI: Vinces, Coastal Lowlands of Ecuador; ECU-PV: Pueblo Viejo, Coastal Lowlands of Ecuador; ECU-QU: Quito, Andean Higlands of Ecuador; ECU-IB: Ibarra, Andean Highlands of Ecuador; GTM-TE: Técpan, Guatemala. GTM-PA: Parramos, Guatemala; GTM-CH: Chimaltenango, Guatemala; GTM-PT: Patzún, Guatemala; GTM-AG: Antigua Guatemala, Guatemala; GTM-CU: Cubulco, Guatemala and GTM-RA: Rabinal, Guatemala. Uncl.: Unclassified species that belong to a taxonomic group but could not be assigned to a specific species. Bars do not reach 100% as species beyond the 30 most abundant are not displayed.

In the Ecuadorian Coastal Lowlands, the relative abundance of *P. maydis* also was low (1.2% to 5.0%). In Mocache and Vinces, *Microdochium seminicola* dominated with 46.3% to 68.1% abundance (Fig. 4). In contrast, samples from Pueblo Viejo had a high abundance of *Paraphaeosphaeria* sp. (42.5%) and unclassified *Dothideomycetes* (25.6%), while *M. seminicola* was almost absent.

In the Andean Highlands of Ecuador, *Phyllachora maydis* and *Paraphaeosphaeria* sp. were predominant, representing nearly 90% of the total abundance. In Ibarra, *P. maydis* represented 26% of the relative abundance, while in Quito, it comprised 50.9% (Fig. 4). Meanwhile, *Paraphaeosphaeria sp.* comprised 59.8% of the relative abundance in Ibarra and 38.9% in the Quito region.

The relative abundance of *Coniothyrium* in Ecuador was less than 0.02%. In the Coastal Lowlands, it was detected in only four out of ten samples; in the Andean Highlands, it was found in seven out of thirteen samples; it was not detected in any of the ten samples collected from the Amazon.

Similar to Ecuador, in Guatemala *Phyllachora maydis*, *Paraphaeosphaeria* sp., and *Microdochium seminicola* were abundant in tar spot lesions (Fig. 4). However, *Aspergillus* spp., particularly *Aspergillus cristatus*, were also prominent. A total of five ASVs were classified as *A. cristatus*, with one of them containing 99.8% of the total sequences assigned to this taxon. This ASV varied by 11 nucleotide positions (96% of similarity) from the representative sequence in the database (Table 3) so it likely represents a different, unnamed species. In Tecpán and Chimaltenango, *Phyllachora maydis* was the predominant species with relative abundances of 71.9% and 26.9%, respectively. The abundance of *Paraphaeosphaeria*, *Microdochium* and *Aspergillus* in these locations was less than 1.3% each. In Parramos and Patzún, *Paraphaeosphaeria* sp. was the predominant species with relative abundances of 63.7% and 38.4%, respectively, followed by *Phyllachora maydis* with relative abundances of 27.1% and 17.9%, respectively. *Aspergillus cristatus* was abundant in Patzún (12.5%) but not in Parramos (0.4%); meanwhile, the relative abundance of *Microdochium* was less than 3.4% in both locations. In Antigua Guatemala, located in Sacatepéquez state, *Phyllachora maydis* was the most abundant species at 68.2%, followed by *Aspergillus cristatus* at 18.6%. The relative abundances of *Microdochium* and *Paraphaeosphaeria* sp. in this location were less than 1%. In Cubulco and Rabinal, located in Baja Verapaz state, *Microdochium* was the predominant species with relative abundances of 17.1% and 31.8%, respectively, followed by *Aspergillus cristatus* with relative abundances of 23.9% and 12.2%, respectively. Relative abundances of *Phyllachora maydis* and *Paraphaeosphaeria* sp. in these locations were less than 4% each.

**Table 3.**
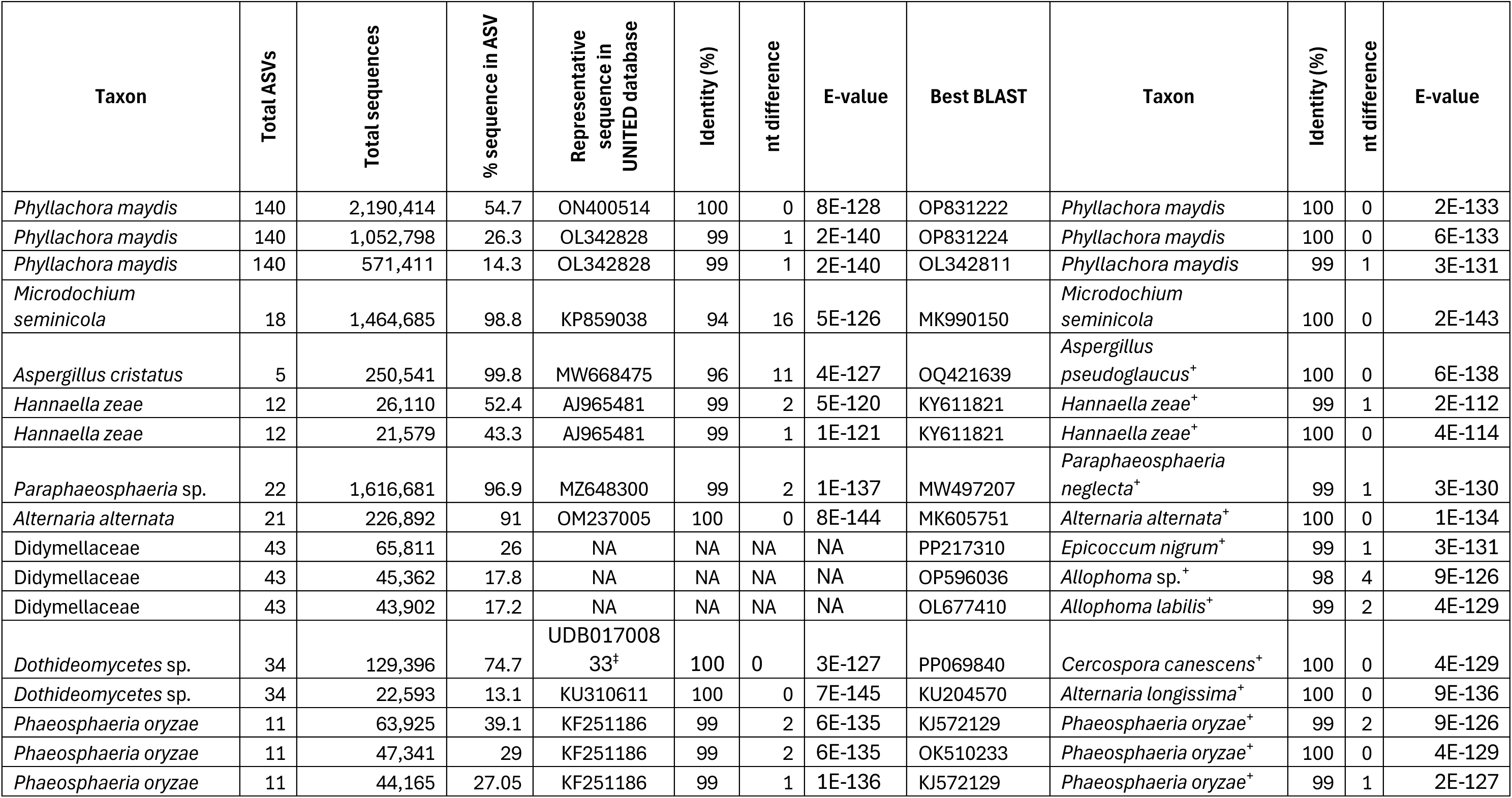
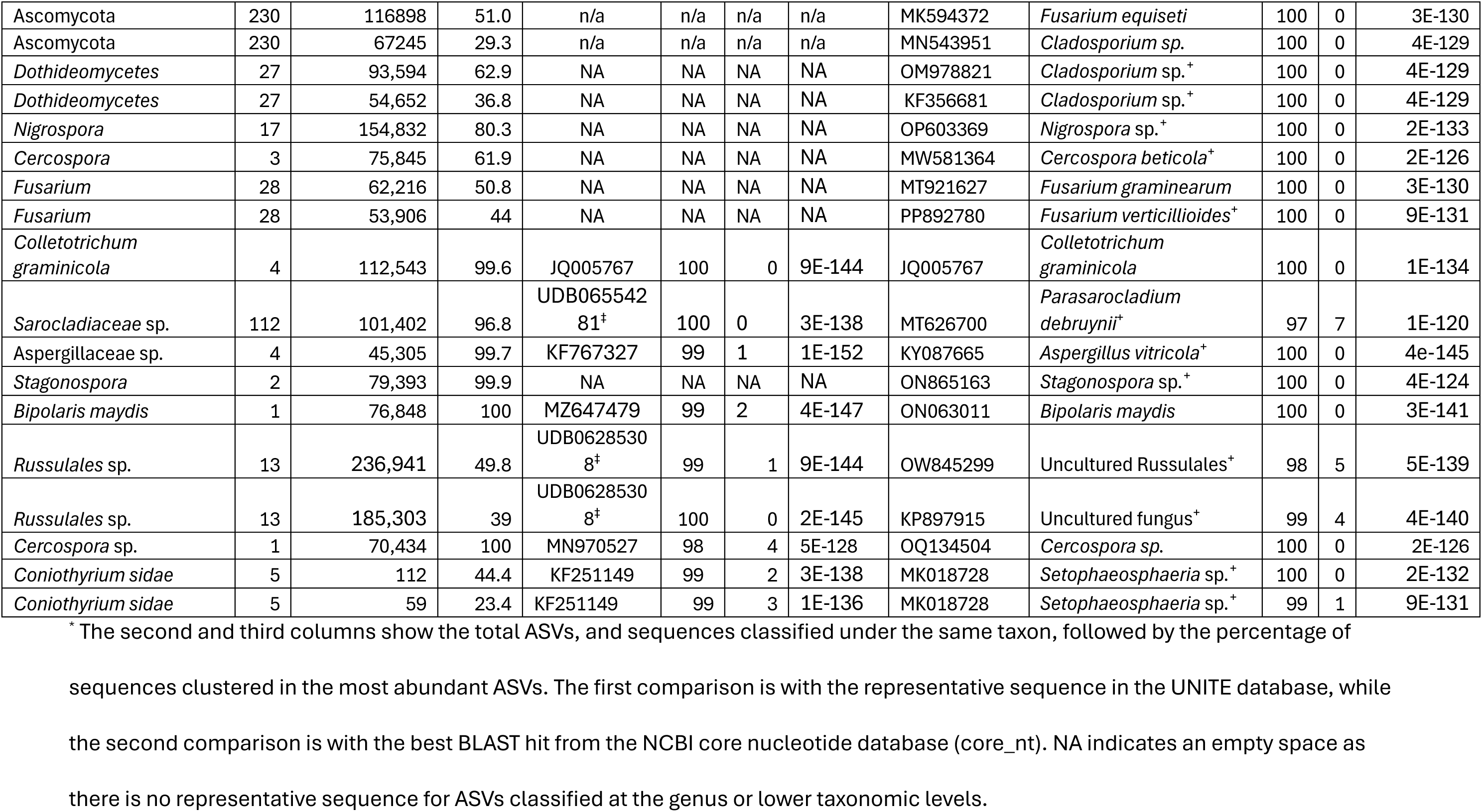
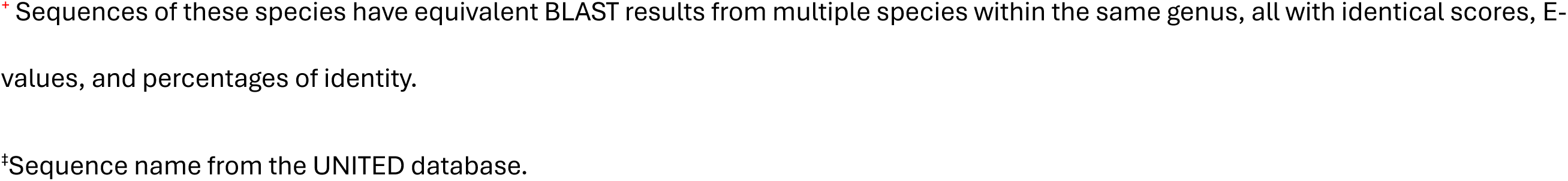
UNITED database and best BLAST results for the most common Amplicon Sequence Variants (ASVs) of the 20 most abundant species plus the possible mycoparasite *Coniothyrium sidae* as the 21^st^ species in tar spot-associated microbiomes*.

Another fungus with variable abundance was the basidiomycetous yeast *Hannaella zeae,* which was the second-most-abundant taxon in Técpan (8.3%) but was almost absent in other locations. A total of 12 ASVs were classified as *H. zeae*, with two of them containing 95.7% of the total sequences assigned to this taxon. These ASVs varied by 1 or 2 nucleotide positions from the representative sequence in the database (Table 3). Unclassified species of the genera *Cercospora* and *Penicilliium* were the second- and third-most-abundant species in Chimaltenango with relative abundances of 16.1% and 10.1% respectively, but they were almost absent in other locations. This tendency also occurs in Rabinal, where the second- and third-most-abundant taxa were unclassified *Fusarium* species and *Aspergillus* sp., with relative abundances of 16.4% and 13.8%, respectively; however, they were almost absent in all other locations, with relative abundances of less than 1%.

*Coniothyrium* was almost absent also in samples collected in Guatemala. We found this species in only five samples from a total of 18 collected in this country and its relative abundance in each of these samples was less than 0.02%. These samples were collected from Técpan, Chimaltenango, Cubulco, and Rabinal.

### *Microdochium seminicola* was predominant in tar spot lesions with fisheye symptoms in Ecuador and Guatemala but not in Indiana, USA

A total of 73 tar spot samples were analyzed across the three countries. Among these, 33 samples did not display fisheye symptoms, whereas 40 samples did. Specifically, 27 fisheye-positive samples were collected from Ecuador, 8 from Guatemala, and 5 from Indiana, USA. The most abundant species in tar spot lesions with fisheye symptoms were *M. seminicola* (22.2% relative abundance), *Paraphaeosphaeria* sp. (20.0%), and *P. maydis* (14.4%). Interestingly, *M. seminicola* was not found in tar spot samples with fisheye symptoms collected in the USA. In contrast, in tar spot lesions with non-fisheye symptoms, *P. maydis* was predominant with a relative abundance of 63.5%, followed by *Paraphaeosphaeria* sp. at 7.1% (Fig. 5).

**Fig 5.**
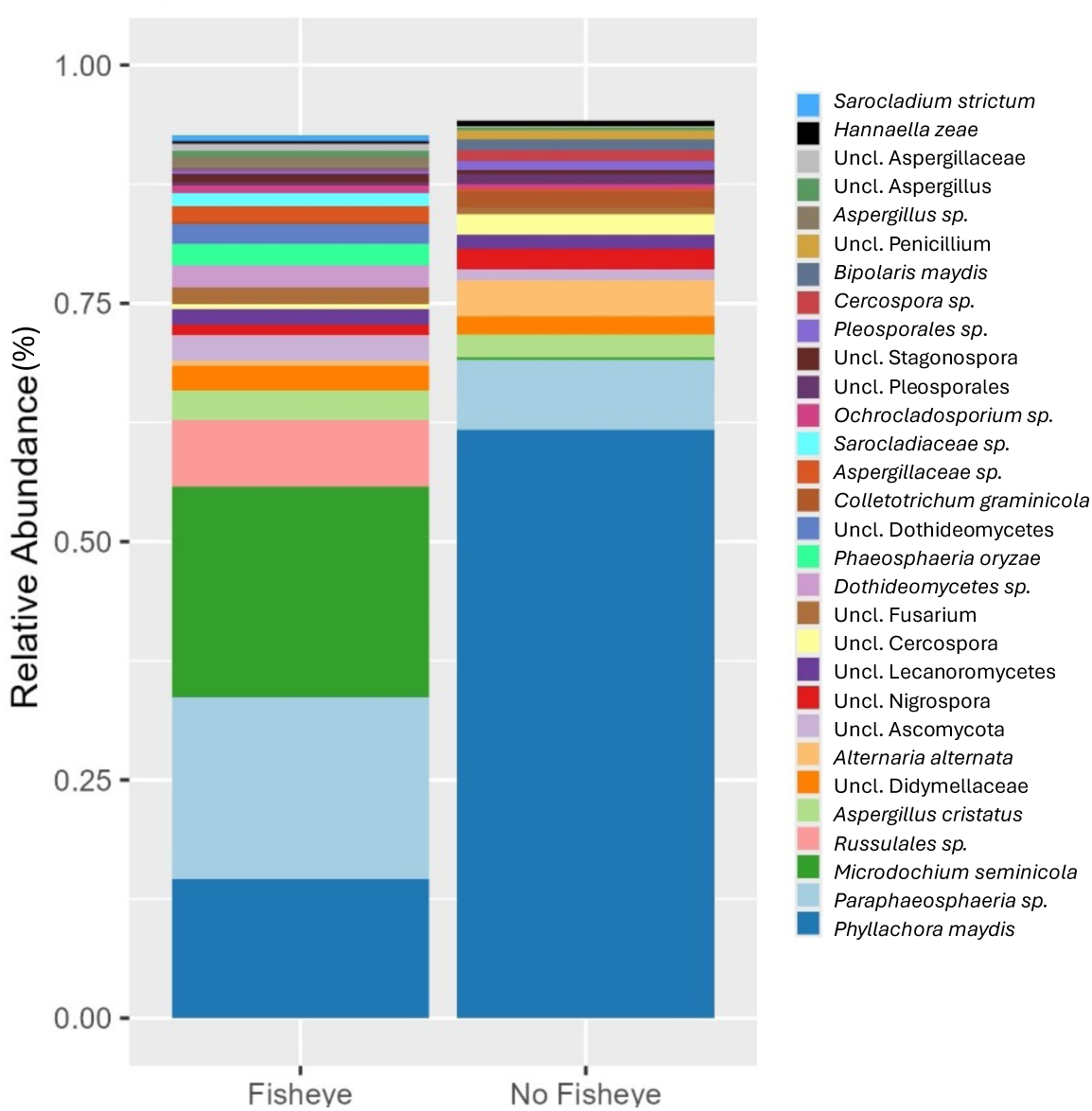
The 30 most abundant fungal species in tar spot lesions with fisheye versus non-fisheye symptoms over all samples. Bars do not reach 100% as species beyond the 30 most abundant are not displayed.

The Analysis of Compositions of Microbes with Bias Correction (ANCOM-BC) revealed 14 genera that were enriched in tar spot lesions with fisheye symptoms compared to those with non-fisheye. These genera include *Microdochium*, *Phaeosphaeria*, *Ochrocladosporium*, *Pyrenochaetopsis*, *Bulleribasidium*, *Strelitziana*, *Curvularia*, *Clonostachys*, *Sarocladium*, *Hannaella*, *Sulcosporium*, a genus in the family *Herpotrichiellaceae*, a genus in the order *Russuales* and another from the class *Dothideomycetes*; the last three genera are classified as genus incertae sedis. In contrast, *Phyllachora*, *Phaeosphaeriopsis*, *Dioszegia*, *Alternaria*, *Symmetrospora*, *Bullera*, *Neosetophoma*, *Didymocyrtis*, *Neoascochyta*, a genus in the family *Mycosphaerellaceae*, one genus in the order *Pleosporales* and another in the phylum *Basidiomycota* were depleted in fisheye lesions, which suggests that these genera are enriched in tar spot lesions with non-fisheye symptoms (Fig. 6). Only *Phyllachora*, *Microdochium*, *Phaeosphaeria*, *Alternaria*, and the genus in the class *Dothideomycetes* had a relative abundance above 1%.

**Fig. 6.**
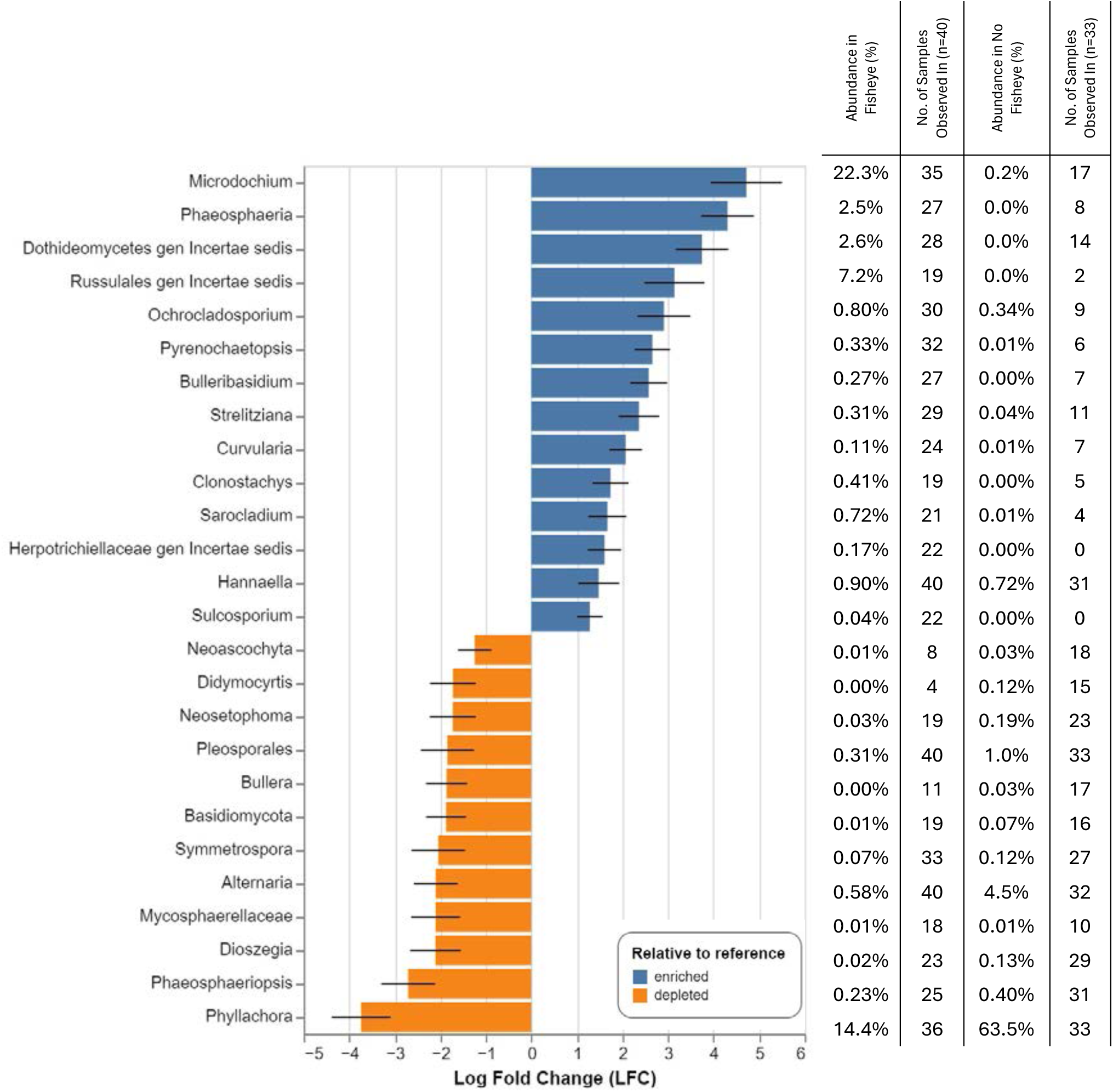
Differentially abundant genera in tar spot samples with fisheye and non-fisheye symptoms, identified using Analysis of Compositions of Microbes with Bias Correction (ANCOM-BC). Blue bars represent genera enriched in fisheye samples compared to those with non-fisheye, while orange bars indicate genera depleted in fisheye lesions, which can also be interpreted as enriched in the non-fisheye samples.

### Tar spot microbiome alpha diversity differs among locations

Samples from the USA exhibited the lowest alpha diversity in terms of both richness and evenness (Fig. 7A, 8A, and 9A). In contrast, alpha diversity in samples from Ecuador and Guatemala varied depending on the specific location and geographic region. Most of the differences were highly significant, whether at the country or region levels (Supplementary Table S1) or in pairwise comparisons (Supplementary Table S2).

**Fig. 7.**
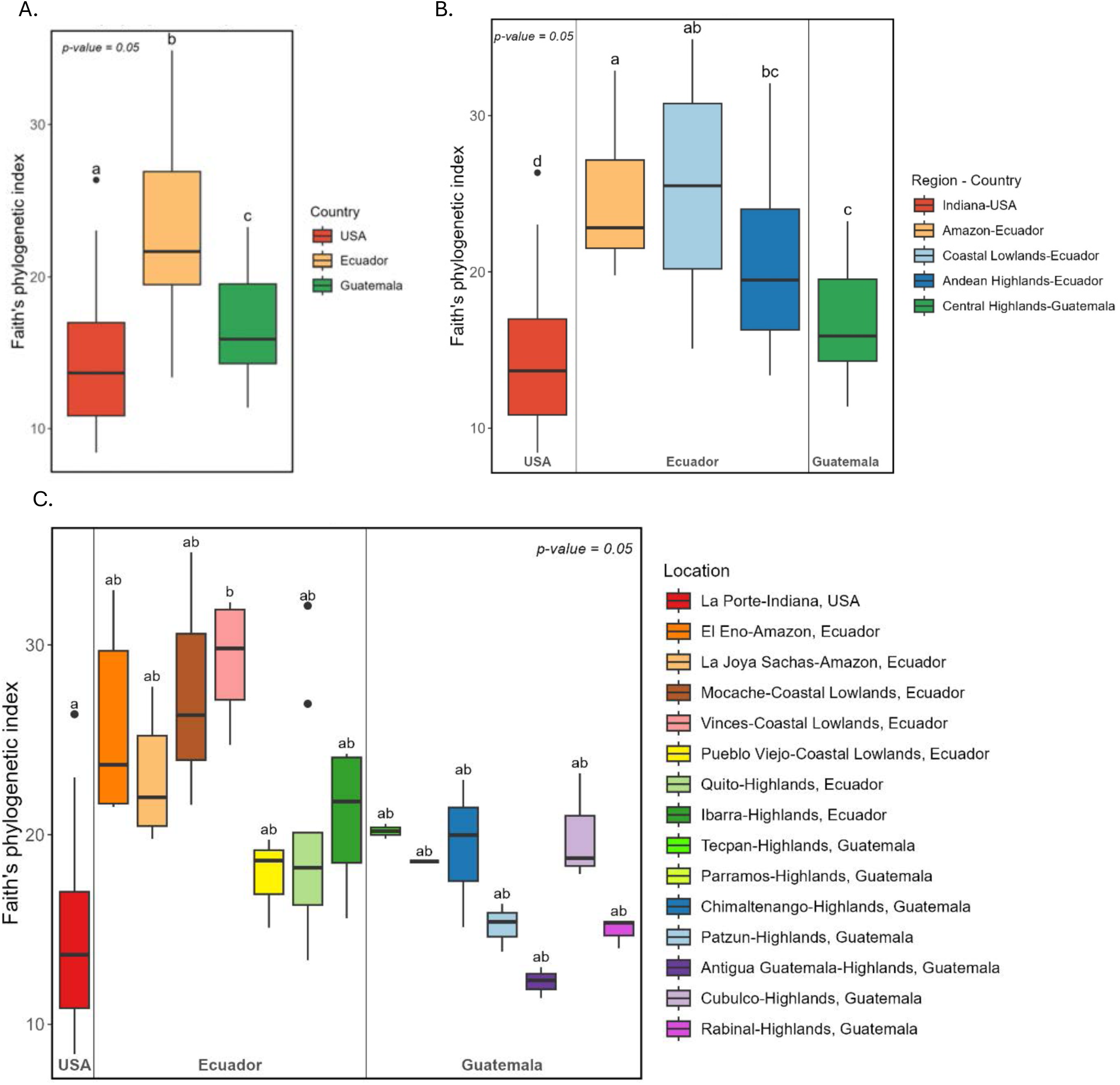
Faith’s phylogenetic index of Alpha diversity among tar spot samples collected in Ecuador, Guatemala and the USA (A), grouped by geographical region (B) and sampling location (C). Bars with different letters show statistical significance at p= 0.05. Whiskers extend to the most extreme data points within 1.5 times the interquartile range (IQR) from the lower and upper quartiles, with points beyond this range plotted as outliers.

**Fig. 8.**
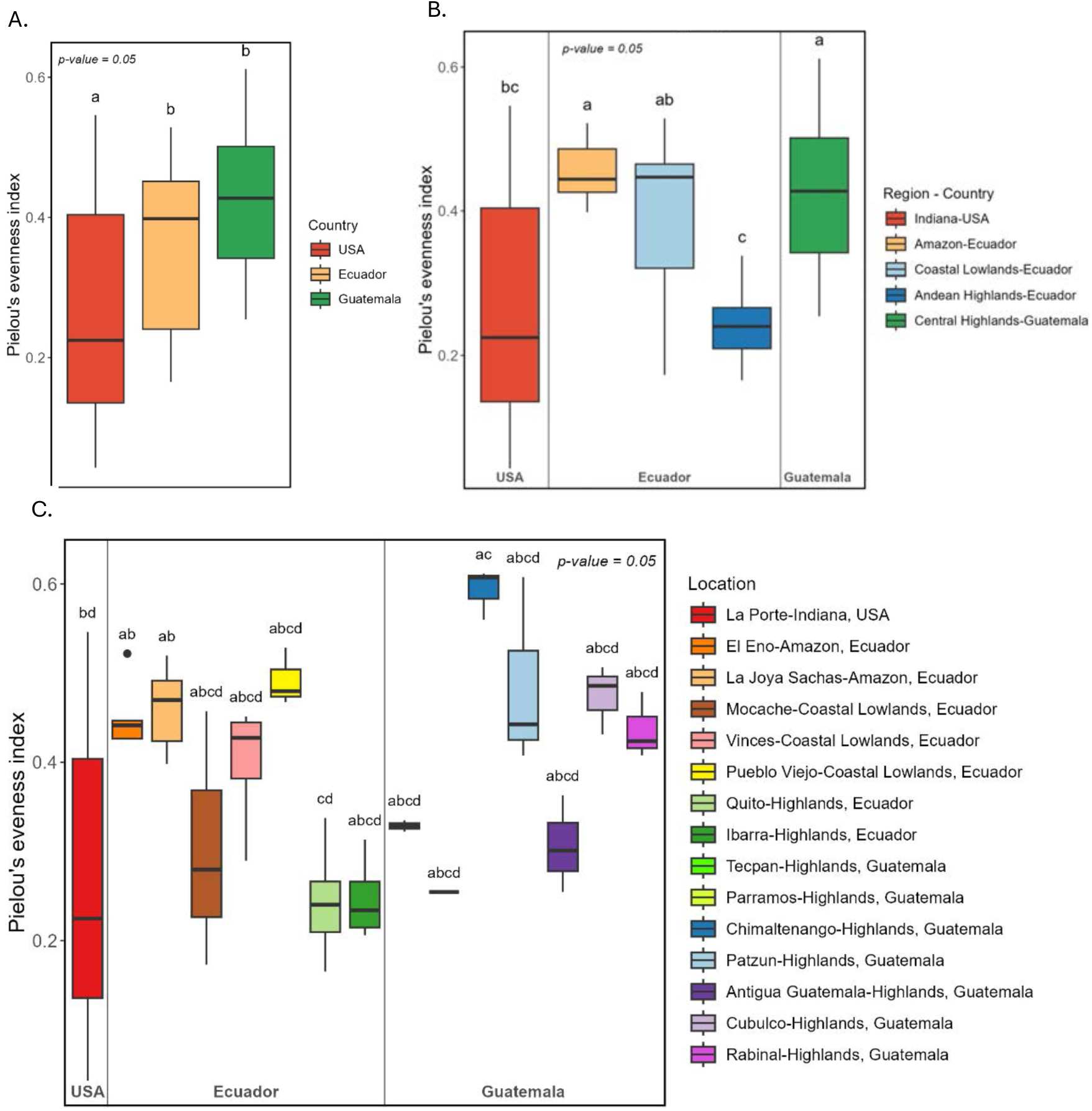
Pielou’s evenness index of tar spot samples collected in Ecuador, Guatemala and the USA (A), grouped by geographical region (B) and sampling location (C). Bars with different letters show statistical significance at p= 0.05. Whiskers extend to the most extreme data points within 1.5 times the interquartile range (IQR) from the lower and upper quartiles, with points beyond this range plotted as outliers.

**Fig. 9.**
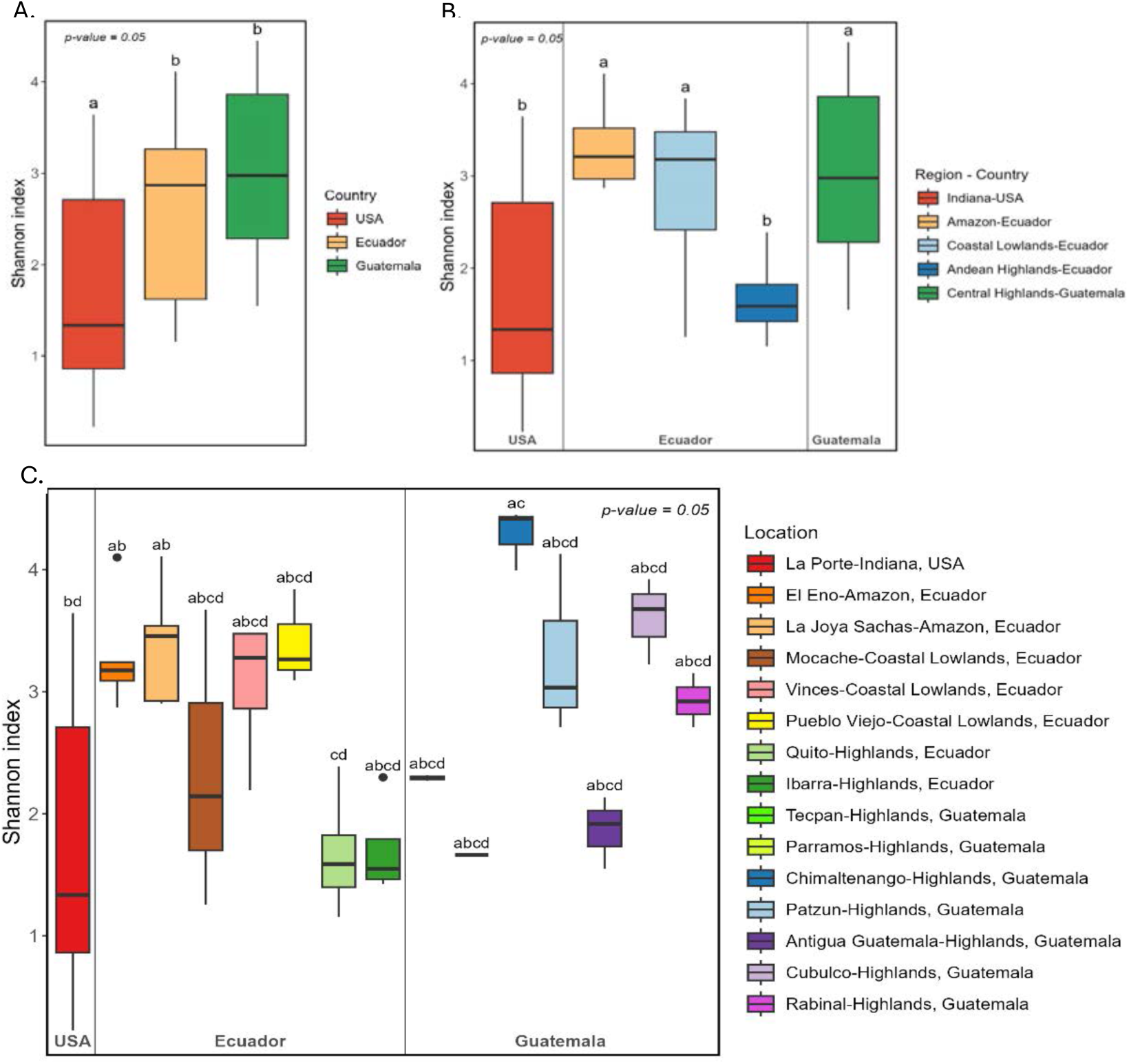
Shannon’s index of Alpha diversity among tar spot samples collected in Ecuador, Guatemala and the USA (A), grouped by geographical region (B) and sampling location (C). Bars with different letters show statistical significance at p= 0.05. Whiskers extend to the most extreme data points within 1.5 times the interquartile range (IQR) from the lower and upper quartiles, with points beyond this range plotted as outliers.

Based on Faith’s phylogenetic index, samples from Indiana exhibited the lowest phylogenetic diversity and were statistically different from those in the geographic regions of Guatemala and Ecuador (Fig. 7B and Supplementary Table S3). In contrast, samples from the Ecuadorian Amazon and Coastal Lowlands of Ecuador had the highest phylogenetic diversity and were statistically similar (Supplementary Table S3). Meanwhile, samples from the highlands of Guatemala and the Andean highlands of Ecuador displayed intermediate diversity and were also statistically similar (Fig. 7B).

Pielou evenness and Shannon indexes grouped the geographical regions into two clusters. The first cluster includes samples from the Ecuadorian Amazon, the Coastal Lowlands of Ecuador, and Central Highlands of Ecuador. The second cluster includes samples from La Porte, Indiana, and the Andean Highlands of Ecuador (Fig. 8B and Fig. 9B).

At location level, the location Vinces, situated in the Ecuadorian lowlands, had the highest richness (Fig. 7C), while Chimaltenango in Guatemala had the highest evenness (Fig. 8C). The lowest richness and evenness were observed in Antigua Guatemala, and Quito, Ecuador, respectively. When both richness and evenness are considered using the Shannon index, Chimaltenango shows the highest diversity while Quito shows the lowest (Fig. 9C). The three measures of alpha diversity were mostly consistent but had some variability across locations, geographic regions, and countries (Figs. 7-9).

The Faith’s phylogenetic index grouped the locations into two clusters (Fig. 7C). The first cluster includes samples from all locations in Guatemala, the Andean Highlands of Ecuador, and La Porte, Indiana. The second cluster includes samples from locations in the Ecuadorian Amazon and the Coastal Lowlands of Ecuador. However, only the samples from La Porte, Indiana, compared to Vinces from the Coastal Lowlands of Ecuador, showed statistical significance. None of the other clusters were statistically different (α= 0.05). When species abundance is considered using Pielou’s evenness index, most locations do not show statistically significant differences. Only the comparisons between the locations Chimaltenango-Guatemala vs. La Porte, USA; Quito-Ecuador vs. El Eno-Ecuador; and Quito-Ecuador vs. La Joya Sachas-Ecuador are statistically different (Fig. 8C). This trend is also observed when both richness and evenness are analyzed together using the Shannon index (Fig. 9C).

### Beta diversity clustered samples into four groups

Based on the Bray-Cutis metric, samples from each country are mostly clustered together (Fig. 10A) and are statistically different from those in other countries (Supplementary Tables S4 and S5). The picture became messier when samples were grouped by geographic region, as some samples from the same region did not cluster together as expected (Fig. 10B). Some samples from one geographic region clustered into another region forming four groups regardless of their geographic region or country (Fig. 11). Each group shared similar fungal species in similar abundances within the group, but these differed between groups. The first group includes all samples from La Porte, Indiana, which do not share any similarities with the other groups. A second group was geographically diverse and clustered all samples from the Ecuadorian Amazon, the locations Vinces and Mocache from the Coastal Lowlands of Ecuador, and the locations Rabinal and Cubulco from Guatemala. Group 3 clustered samples from all locations in the Andean Highlands of Ecuador, the location Pueblo Viejo from the Coastal Lowlands of Ecuador, Parramos from Guatemala, and two samples from Patzún, Guatemala. A third sample from Patzún, Guatemala clustered with samples from Técpan, Chimaltenango, and Antigua Guatemala to form the fourth group. Pairwise comparisons between geographic regions revealed statistically significant differences (Supplementary Table S6) for all groups, except for the Ecuadorian Amazon and the Coastal Lowlands, which were not statistically different and were grouped in cluster 2.

**Fig. 10.**
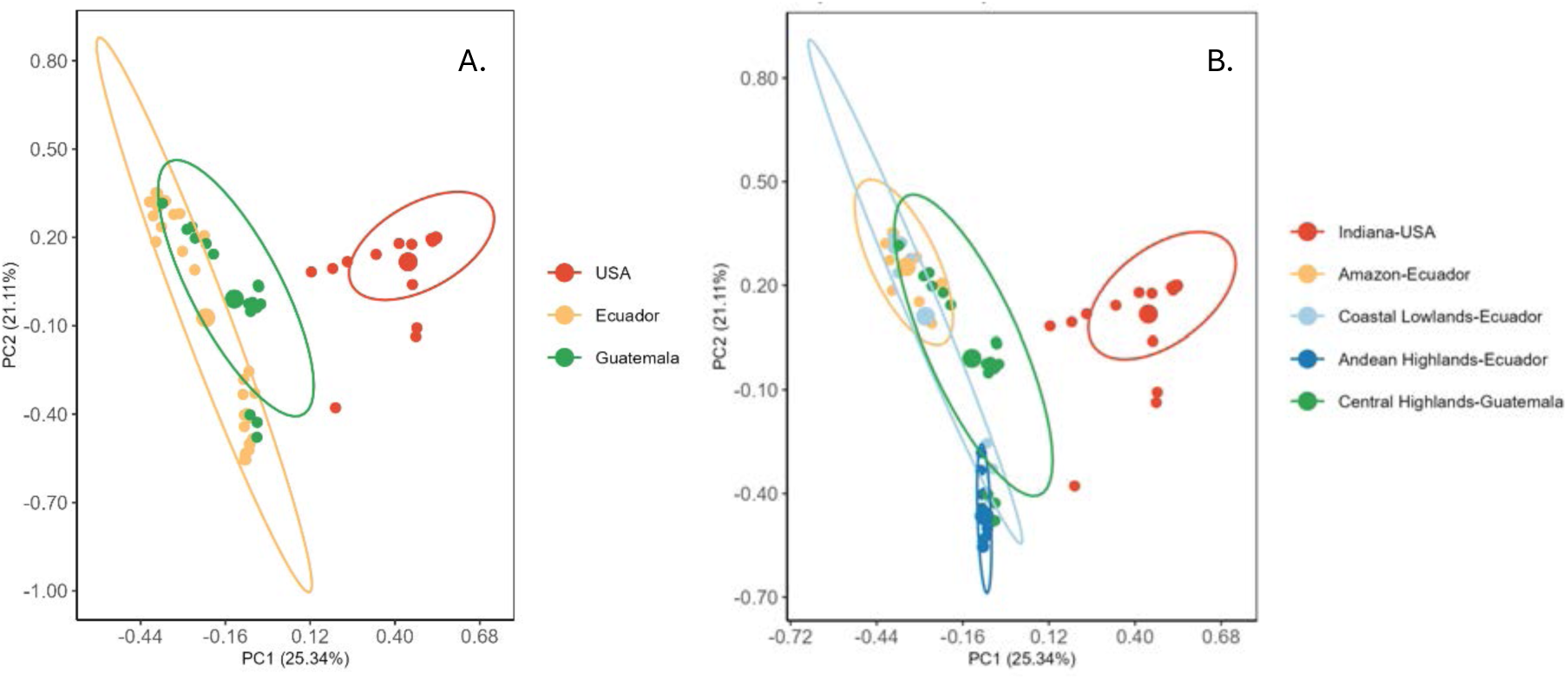
Principal coordinate analysis (PCoA) of Bray–Curtis dissimilarity between countries (A) and geographic regions (B). Ellipses indicate the 95% confidence interval of samples belonging to each group. Larger circle dots indicate the centroids of the ellipses.

**Fig. 11.**
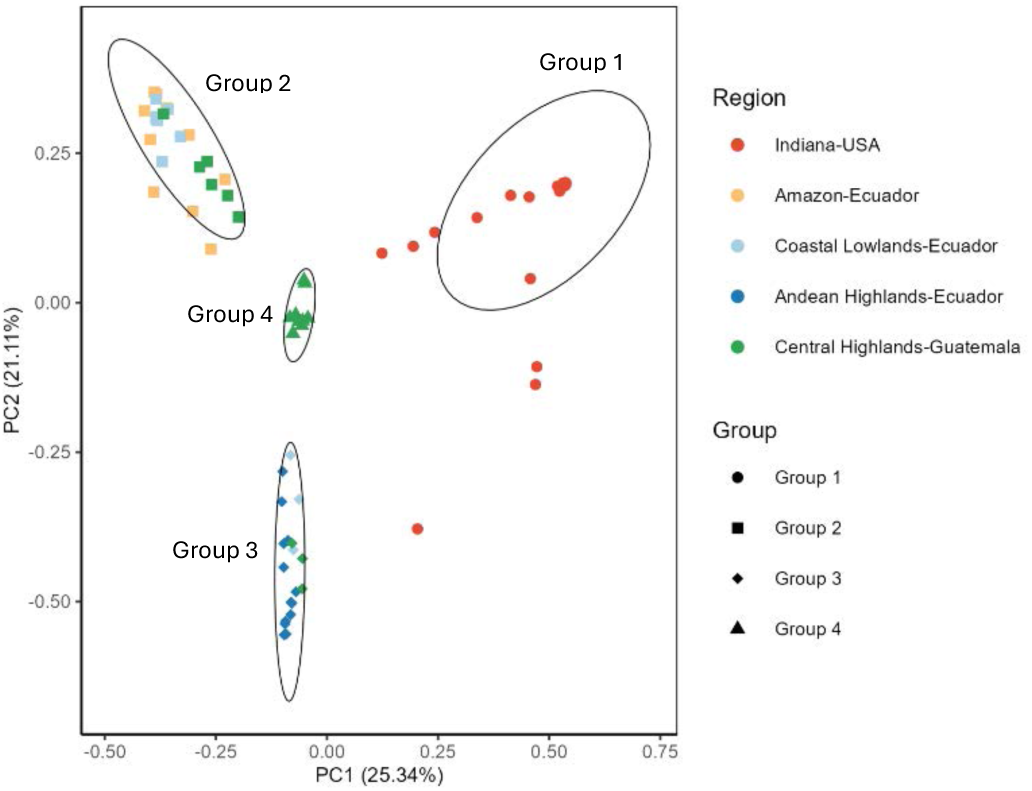
Principal coordinate analysis (PCoA) of Bray-Curtis dissimilarity between groups. Each group consists of tar spot samples with a predominant species. In groups 1 and 4 the predominant species is *Phyllachora maydis*; in the second group the predominant species is *Microdochium seminicola*; and in the third group *Phyllachora maydis* and *Paraphaeosphaeria* sp. were the predominant species. Ellipses circumscribe the 95% confidence interval of the samples belonging to each group.

Samples from each country and geographic region formed less distinct clusters based on the weighted UniFrac metric (Fig 12). A clear separation was observed only between samples from the Ecuadorian Amazon and those from Indiana, as well as between the Amazon and the Andean highlands of Ecuador. In contrast, samples from Indiana, the Andean highlands of Ecuador, the highlands of Guatemala, and the Coastal Lowlands of Ecuador showed significant overlap. The pairwise comparison between countries and geographic regions revealed statistically significant differences (Supplementary Tables S5 and S6) for all regions, except for the comparison between the Ecuadorian Amazon and the Coastal Lowlands, which did not show a statistically significant difference. This finding is consistent with the results obtained from pairwise comparisons using the Bray-Curtis index.

**Fig. 12.**
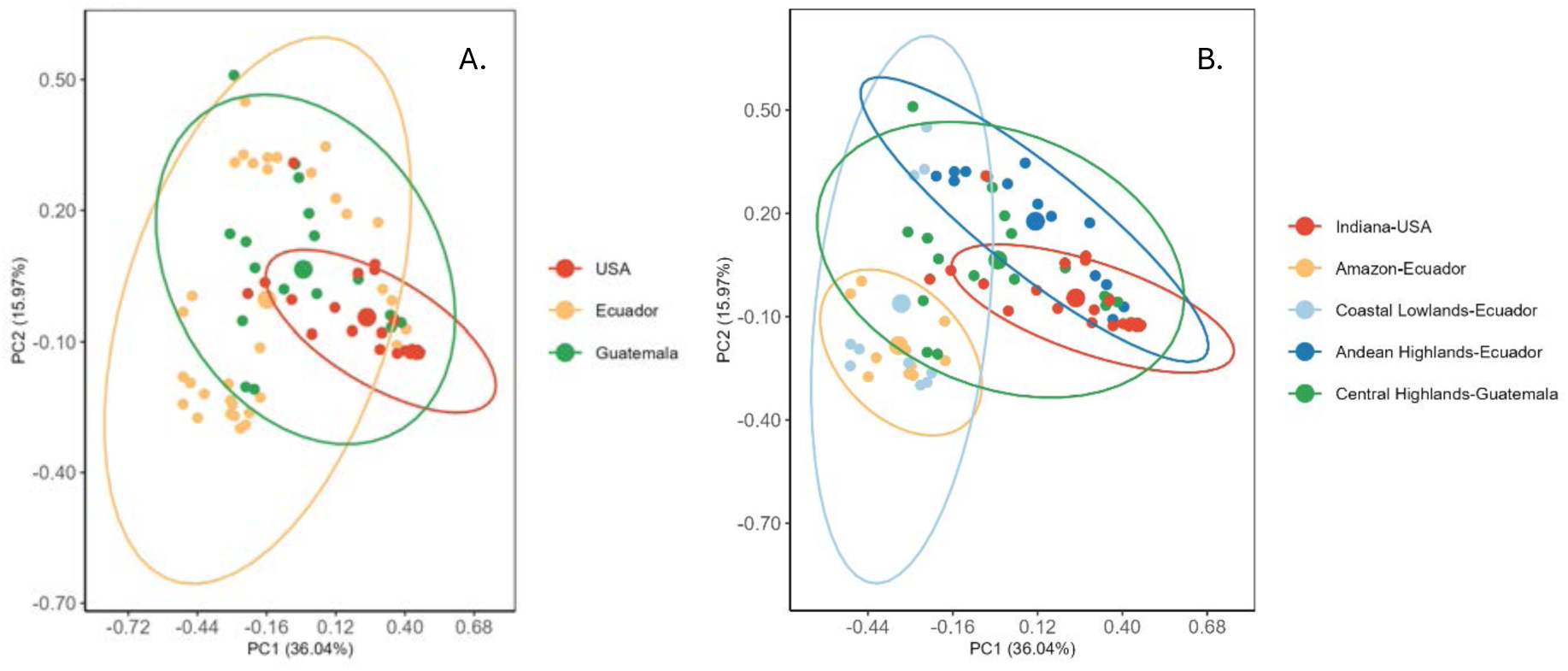
Principal coordinate analysis (PCoA) of Weighted UniFrac beta diversity between countries (A) and geographic regions (B). Ellipses indicate the 95% confidence interval of samples belonging to each group. Larger circle dots indicate the centroids of the ellipses.

### Correlations between the most abundant species

We tested associations among the twenty most abundant fungal species in tar spot lesions across all samples. Due to its possible role as a mycoparasite *Coniothyrium sidae* was included as the 21^st^ species despite its low abundance (Fig. 13). There were 247 positive correlations and 194 negative correlations. The strongest correlations with *Phyllachora maydis* were with *Phaeosphaeria oryzae* (Spearman correlation coefficient, ρ = −0.73), *Microdochium seminicola* (ρ = −0.72), a species of the class *Dothideomycetes* (ρ = −0.67), and *Alternaria alternata* (ρ = 0.52), all showing statistical significance at p = 0.05 (Table 4). The correlation coefficients between *P. maydis* with *Paraphaeosphaeria* sp. and *Coniothyrium sidae* were 0.09 and −0.12, respectively, but neither was statistically significant.

**Fig. 13.**
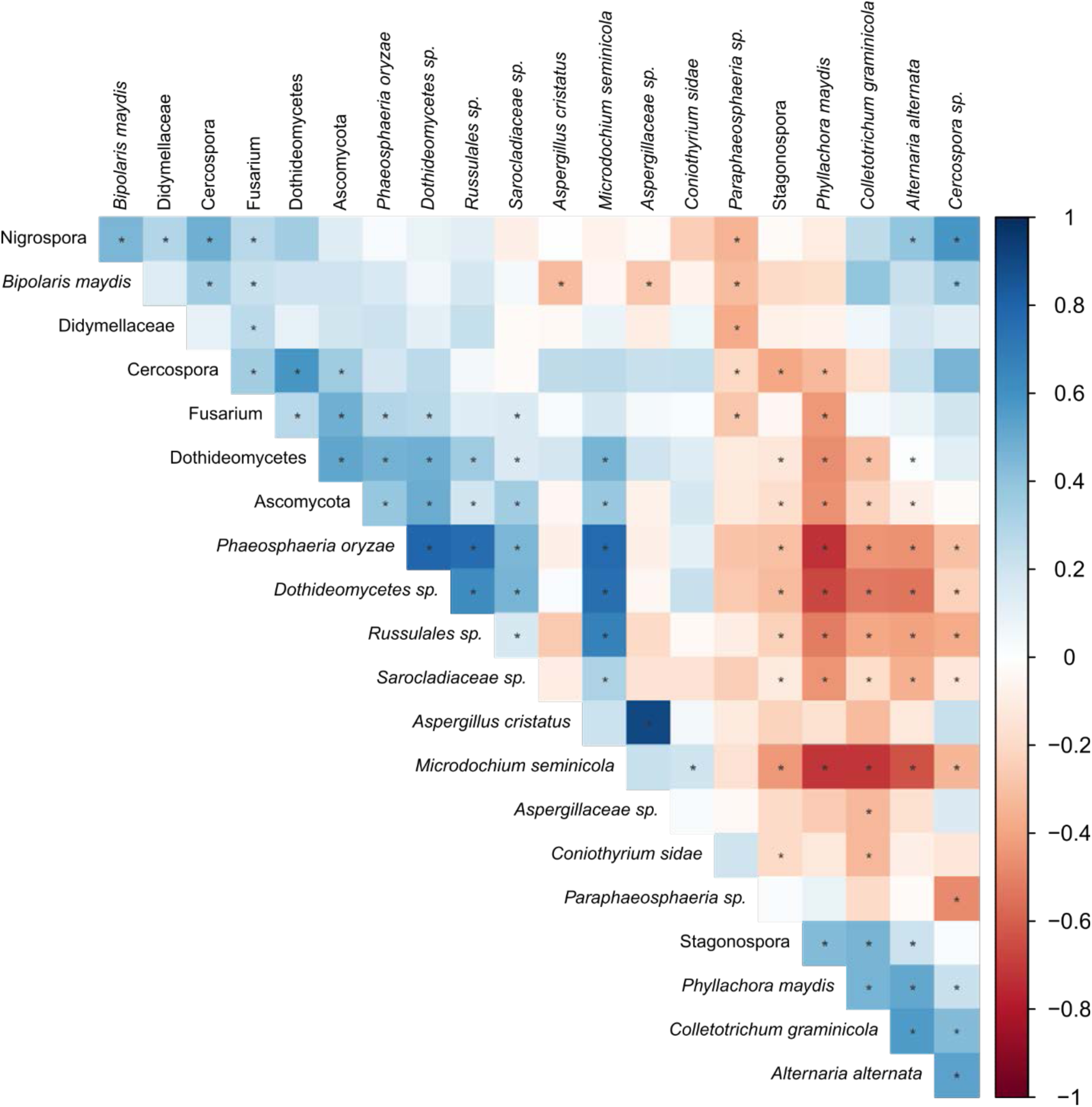
Heat map of the Spearman correlation coefficients for all pairwise correlations between the 20 most-abundant fungal species in tar spot lesions. *C. sidae* was included as the 21^st^ species. * Statistical significance at p = 0.05.

**Table 4.** Spearman’s rank correlation coefficients between *Phyllachora maydis*, *Paraphaeosphaeria* sp., *Microdochium seminicola* and *Coniothyrium sidae* with the 20 most abundant species.

| Species | *Phyllachora maydis* | *Paraphaeosphaeria* sp. | *Microdochium seminicola* | *Coniothyrium sidae* |
| --- | --- | --- | --- | --- |
| *Phyllachora maydis* | 1.00 | 0.09 | -0.72*** | -0.12 |
| *Paraphaeosphaeria* sp. | 0.09 | 1.00 | -0.15 | 0.20 |
| *Microdochium seminicola* | -0.72*** | -0.15 | 1.00 | 0.20* |
| Russulales order | -0.52*** | -0.10 | 0.67*** | -0.04 |
| Didymellaceae family | -0.06 | -0.38** | 0.09 | 0.07 |
| *Aspergillus cristatus* | -0.16 | -0.11 | 0.21 | 0.05 |
| *Alternaria alternata* | 0.52*** | -0.02 | -0.64*** | -0.09 |
| Ascomycota | -0.45*** | -0.13 | 0.37*** | 0.17 |
| *Nigrospora* genus | -0.10 | -0.34** | -0.06 | -0.24 |
| *Dothideomycetes* sp. | -0.67*** | -0.26 | 0.75*** | 0.23 |
| *Phaeosphaeria oryzae* | -0.73*** | -0.29 | 0.77*** | 0.11 |
| *Dothideomycetes* | -0.47*** | -0.11 | 0.47*** | 0.14 |
| *Cercospora* genus | -0.33* | -0.21* | 0.26 | 0.23 |
| *Fusarium* genus | -0.44** | -0.28** | 0.18 | 0.03 |
| *Colletotrichum graminicola* | 0.46*** | -0.20 | -0.71*** | -0.33** |
| *Sarocladiaceae* sp. | -0.44*** | -0.24 | 0.31*** | -0.16 |
| *Aspergillaceae* sp. | -0.25 | -0.03 | 0.23 | 0.04 |
| *Stagonospora* genus | 0.44*** | 0.02 | -0.44*** | -0.20* |
| *Bipolaris maydis* | -0.17 | -0.32** | -0.04 | -0.07 |
| *Cercospora* sp. | 0.22* | -0.48* | -0.33** | -0.14 |
| *Coniothyrium sidae* | -0.12 | 0.20 | 0.20* | 1.00 |

Asterisks indicate significant p values at $\leq 0.05$ (\*), $\leq 0.01$ (\*\*), and $\leq 0.001$ (\*\*\*).

*Microdochium seminicola* was negatively correlated with *Alternaria alternata* (ρ = −0.64) and *Colletotrichum graminicola* (ρ = −0.71), but positively associated with *Phaeosphaeria oryzae* (ρ = 0.77), one species of the Dothideomycetes class (ρ = 0.75), and one species from the order Russulales (ρ = 0.67), all showing statical significance at p = 0.05 (Table 4).

Most correlations involving *Paraphaeosphaeria* sp. were not statistically significant at α = 0.05 (Table 4). Among the statistically significant correlations, all were negative, including those with a species from the family Didymellaceae (ρ = −0.38), and the genera *Fusarium* (ρ = −0.28) and *Nigrospora* (ρ = −0.34). Similarly, most correlations with *Coniothyrium sidae* were also not statistically significant at p = 0.05 (Table 4). Notably, *C. sidae* exhibited a positive correlation with *Microdochium seminicola* (ρ = 0.20) and negative correlations with *Colletotrichum graminicola* (ρ = −0.33), and a species from the genus *Stagonospora* (ρ = −0.20).

## DISCUSSION

*Phyllachora maydis* and *Coniothyrium phyllachorae* were described as new in 1904 (Maublanc, 1904). In his report, Maublanc also stated, “this species was accompanied by a fungus which in Mexico causes considerable damage to young corn plants, and which does not seem to me to differ from *Helminthosporium turcicum* Pass., already reported as a parasite of corn”. It is possible that he was describing *Microdochium maydis* (syn. *Monographella maydis*) causing fisheye lesions. However, it was not until 1984 that *Monographella maydis* was described as a new species in tar spot disease from samples collected in Poza Rica, Mexico (Muller & Samuels, 1984). By 1991 tar spot was already considered as a complex (Hock et al., 1992). Morphological identification and fungal isolation have been conducted in Guerrero (Pereyda-Hernández, 2009) and Chiapas, Mexico and in Guatemala (Sol Hernandez, 2018) showing tar spot as a complex of three fungi with *P. maydis* as the primary pathogen, *M. maydis* possibly causing fisheye lesions and *Coniothyrium phyllachorae* as a possible mycoparasite of *P. maydis*. Which species occur where, and exactly how they interact are not known.

In the USA, tar spot was not detected until 2015 (Ruhl et. al., 2016) and since then it has been spreading in the northern states as far west as Nebraska and the Dakotas, and in the south to Georgia and Florida. In the USA, tar spot has been associated only with *Phyllachora maydis*, since *Monographella maydis* has not yet been identified and *Coniothyrium* has been found in low abundance. In fact, Luis et al. (2023) did not identify *Microdochium* in fungi isolated from tar spot lesions with and without fisheye symptoms from samples collected in Florida, Illinois, and Wisconsin in the USA and Chiapas, Guerrero, Oaxaca, Puebla, and Veracruz in Mexico. They isolated 96 *Microdochium*-like fungi and concluded that they belong to the genus *Fusarium*. Furthermore, a microbiome analysis conducted by McCoy et al. (2019) with samples collected in Michigan found only two OTUs identified as *Microdochium* in 12 samples with a maximum number of reads of eight, concluding that *Monographella maydis* is not required for fisheye development in Michigan. They did not report *Coniothyrium*; however, they did report a high abundance of *Paraphaeosphaeria* in tar spot lesions with fisheye symptoms. A limitation for identifying these species is the availability of reference sequences since neither *C. phyllachorae* nor *M. maydis* have been isolated.

Is tar spot of corn caused by a complex of three fungal species? Could the existence of *Microdochium* and *Coniothyrium* in Mexico, Central, and South America have been misidentified? The result of our research indicates that *P. maydis* is not the sole fungus present in tar spot lesions. Instead, these lesions can consist of a combination of two to three different fungal species. For instance, in Indiana, USA, *P*. *maydis* and *Paraphaeosphaeria* sp. are highly abundant in tar spot lesions, while in Mocache, Ecuador, the lesions are dominated by *P. maydis* and *Microdochium seminicola*. In Patzún, Guatemala, a combination of *P. maydis*, *Paraphaeosphaeria* sp., and *Aspergillus* spp. is observed. These findings suggest that tar spot microbiomes are geographically variable and frequently involve multiple fungal species, including *P. maydis*, *Paraphaeosphaeria* sp., and *M. seminicola*. The roles of *Paraphaeosphaeria* and *Microdochium* in tar spot remain unclear. For instance, *M. maydis* was associated with fisheye lesions in some samples from Ecuador and Guatemala, yet its near absence in the USA raises questions about its role as a primary pathogen or opportunist.

*Coniothyrium* was detected at low abundance, comprising less than 1%, in only 15 out of 73 samples collected across the three countries. In contrast, we observed a high abundance of *Paraphaeosphaeria*, a member of the order Pleosporales, in most of the samples. These findings differ from those of Singh et al. (2024, unpublished data), who reported a high abundance of *Coniothyrium* in resistant corn lines from Indiana. This discrepancy may be due to differences in database versions, as the UNITED database version 9.0 classified these sequences as *Coniothyrium*, while version 10.0 classifies them only at the level of order, such as Pleosporales. *Coniothyrium*-like fungi are Coelomycetous asexual morphs of the order Pleosporales and other Dothideomycetes (Verkley et al., 2014). The form class Coelomycetes is defined by the morphological characteristics of its asexual reproductive structures. However, molecular studies indicate that the taxonomy of this class is artificial and distributed its members into three classes of the phylum Ascomycota: Dothideomycetes, Leotiomycetes, and Sordariomycetes (Valenzuela-Lopez et al., 2017). Hundreds of species have been described as *Coniothyrium* morphologically based on material found on plants, and their taxonomy is based mostly on their hosts (Verkley et al., 2004); as a result this genus is considered polyphyletic. The taxonomy of *Coniothyrium*-like fungi has been re-assessed using molecular phylogenetic tools (Verkley et al., 2004, 2014), placing many species into the genus *Paraphaeosphaeria*. Based on the first report of *Coniothyrium phyllachorae*, which was identified by morphology, we suspect that this *Coniothyrium*-like species in the tar spot microbiome may be a member of the genus *Paraphaeosphaeria.* Furthermore, Caldwell et al. (2023) identified fungal structures within *P. maydis* stromata; ITS sequencing and morphological analyses suggested that these structures likely were produced by species of *Paraphaeosphaeria*. Therefore, a reassessment of the taxonomy of *Coniothyrium phyllachorae* is required to fully understand which species are associated with tar spot disease in the field.

The abundance of *Paraphaeosphaeria* in tar spot lesions is notably higher in lesions exhibiting fisheye symptoms. McCoy et al. (2019) reported a relative abundance of *Paraphaeosphaeria* of 18.1% in fisheye lesions, compared to 3% in lesions containing only tar spot. The findings of this study align with this, as we observed a relative abundance of *Paraphaeosphaeria* sp. of 37.1% in fisheye lesions, and just 0.03% in non-fisheye lesions from samples collected in Indiana. Samples with fisheye lesions were obtained in 2023, while those with non-fisheye symptoms were collected in both 2021 and 2023. Additionally, the presence and abundance of fisheye lesions was higher in 2021 than 2023 in LaPorte, Indiana, where these samples were collected (personal observation).

The abundance of *Paraphaeosphaeria* sp. was not consistently high in all lesions exhibiting fisheye symptoms. In Ecuador, tar spot lesions can be categorized into three types based on their appearance: lesions with a thick stroma and no fisheye, lesions with a thick stroma surrounded by fisheye, and lesions with a small to medium stroma surrounded by fisheye (Fig. 14). Lesions with thick stromata and no fisheye were found exclusively in Quito, located in the Andean highlands, whereas lesions exhibiting fisheye symptoms were observed in all other locations. Tar spot lesions with thick stromata, either with or without-fisheye symptoms (Figs. 13A and 13B), exhibited a high abundance of *Paraphaeosphaeria* sp. Lesions with thick stromata and no fisheye were found in samples from Quito, Andean Highlands, while lesions with thick stromata surrounded by fisheye were observed in Pueblo Viejo in the Coastal Lowlands and Ibarra in the Andean Highlands. In these locations, *Paraphaeosphaeria* sp. was more abundant than *P. maydis*. In contrast, lesions with small to medium stromata surrounded by fisheye (Fig. 14C) exhibited a low abundance of *Paraphaeosphaeria* sp. at less than 0.1%. This type of lesion was observed in El Eno and La Joya Sachas from the Amazon region, as well as Mocache and Vinces in the Coastal Lowlands of Ecuador.

**Fig. 14.**
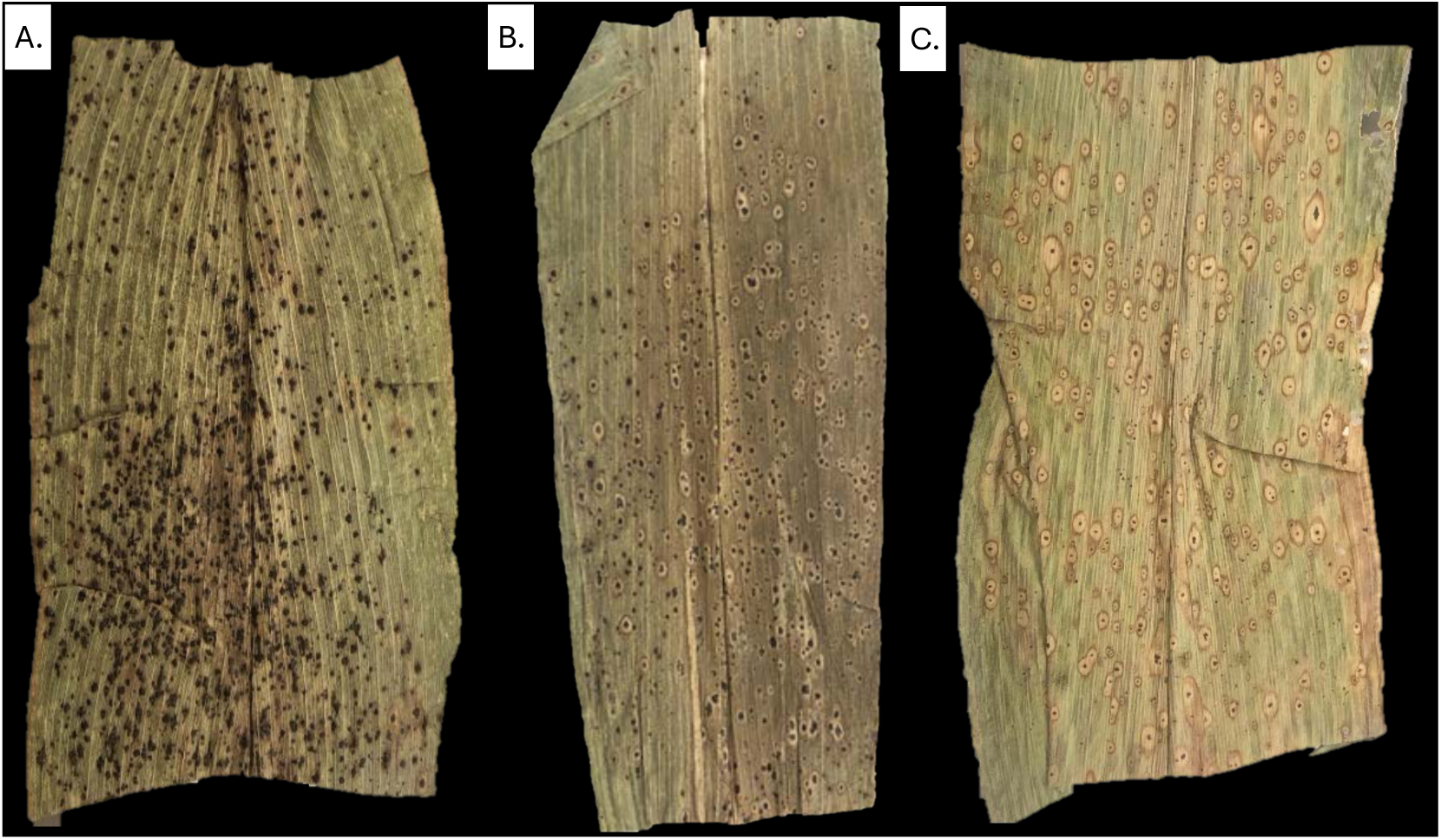
Three types of tar spot lesion found in Ecuador. A. Thick stromata with non-fisheye symptoms, sample collected in Quito, Andean Highlands of Ecuador. B. Thick stromata with fisheye symptoms, sample collected in Ibarra, Andean Highlands. C. Small/medium stromata with fisheye symptoms, sample collected in Vinces, Coastal Lowlands of Ecuador.

In Guatemala, tar spot lesions are categorized only into lesions with thick stromata and no fisheye and lesions with small stromata surrounded by fisheye (Fig. 15). *Paraphaeosphaeria* sp. was present in both tar spot lesions with and without fisheye symptoms. A high abundance of *Paraphaeosphaeria* sp., exceeding that of *P. maydis*, was observed in samples from Parramos, which had stromata without fisheye, and in Patzún, where stromata with fisheye lesions were present. Similar to the samples from Ecuador, we detected a high abundance of *Paraphaeosphaeria* sp. in tar spot lesions with thick stromata and fisheye in Guatemala. However, thick stromata with non-fisheye symptoms did not consistently show a high abundance of *Paraphaeosphaeria* sp. in Guatemala. In fact, we observed only one sample with thick stromata and fisheye that had a high abundance of *Paraphaeosphaeria* sp.

**Fig. 15.**
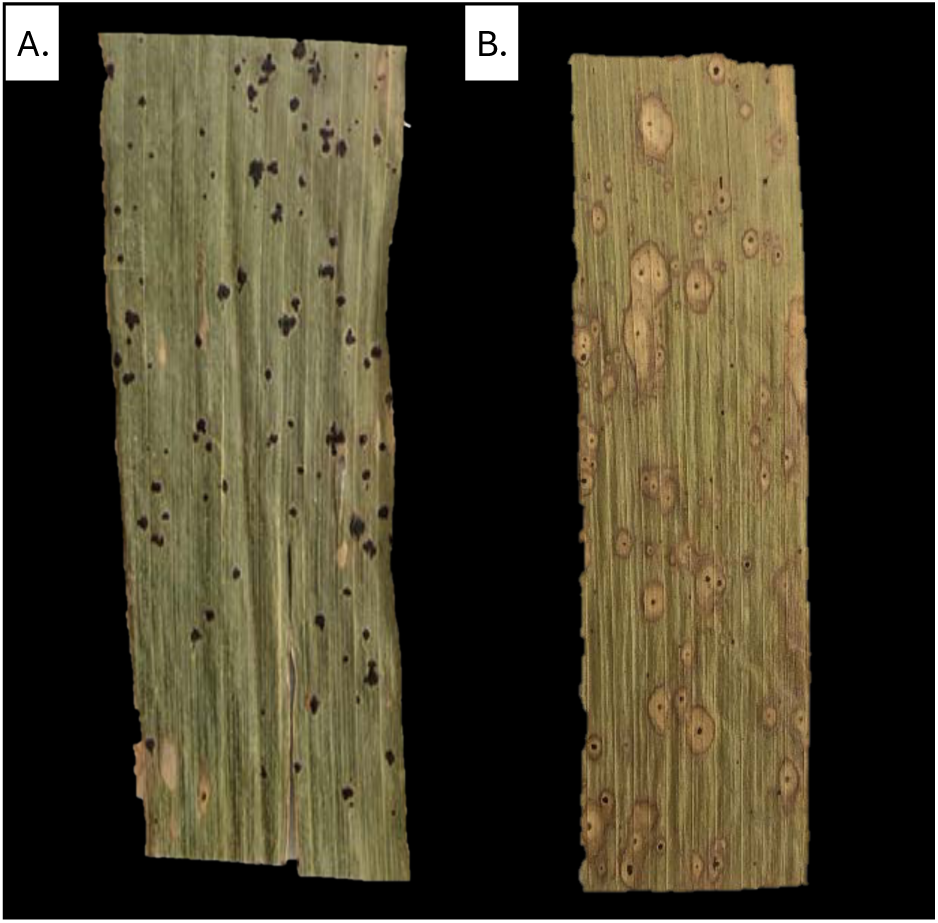
Types of tar spot lesion found in Guatemala. A. Thick stromata with non-fisheye symptoms, sample collected in Antigua Guatemala, Sacatepéquez. B. Small stromata with fisheye symptoms, sample collected in Cubulco, Baja Verapaz.

A high abundance of *Paraphaeosphaeria* sp. was observed in tar spot lesions with fisheye symptoms, but also in thick and small stromata without fisheye. Therefore, the results of this study suggest that the presence of fisheye lesions and stroma size, and the high abundance of *Paraphaeosphaeria* sp. in Ecuador and Guatemala seems to follow a geographical distribution.

*Microdochium maydis* is considered the anamorph of *Monographella maydis* (Muller & Samuels, 1984). However, with the changes implemented by the International Code of Nomenclature for algae, fungi and plants (ICN) (Rossman, 2014) fungal species may have only one name. As a result, *Monographella maydis* is now known as *Microdochium maydis* (Hernández-Restrepo et al., 2016). So far, there are no sequences of *Microdochium maydis* available in the NCBI database despite reports of its isolation from Mexico and Guatemala. Sol Hernandez (2018) isolated strains with characteristics of *Microdochium maydis*, whose ITS sequences showed similarity to *Monographella* sp. and *Microdochium seminicola* species from the NCBI. She also extracted DNA from fisheye lesions, and the ITS sequence also matched with *Monographella* sp. and *Microdochium seminicola*. However, her sequences were not submitted to the NCBI so we cannot confirm how close their sequences are to those from Ecuador and Guatemala.

*Microdochium maydis* was described as the causal agent of fisheye lesions (Hock et al., 1992), with high abundance observed at the late stage of the disease. However, not all tar spot stromata are surrounded by fisheye lesions, and in some locations, such as the USA, their frequency is low (Rocco da Silva et al., 2021). Attempts to confirm the presence of *Microdochium* spp. in tar spot lesions in the USA have been unsuccessful. A culture-based study recovered 101 *Microdochium*-like isolates from necrotic lesions surrounding tar spot, but over 90% of these were identified as *Fusarium* spp. Other isolates were identified as *Epicoccum nigrum*, *E. sorghinum*, or *Irpex lacteus*. These isolates were collected from Florida, Illinois, Wisconsin, and Mexico (Luis et al., 2023). The authors suggested that the initial report of *M. maydis* may have been a misidentification of a *Fusarium* sp. They also suggested that *Fusarium* in tar spot colonizes necrotic tissue rather than being the primary cause of fisheye lesions. Furthermore, the microbiome analysis of tar spot conducted in Michigan supports the hypothesis that *Monographella maydis* is not required for the development of fisheye lesions (McCoy et al., 2019).

Regardless of the hypothesis that *Microdochium maydis* causes fisheye lesions, the result of this study show that *Microdochium* is present at a high frequency in some tar spot lesions in Ecuador and Guatemala but is almost absent in the USA. More than 99% of our *Microdochium* sequences had their highest match with *Microdochium seminicola*. This species was described from isolates found in harvested grains, including oat, barley, and wheat collected in Canada and Switzerland. The description of *Microdochium seminicola* noted that an isolate obtained from maize kernels in Switzerland resembles *Microdochium maydis*, but *M. maydis* conidia are smaller with more septa (Hernández-Restrepo et al., 2016). Our *Microdochium seminicola* sequences were clustered into 18 Amplicon Sequence Variants (ASV) but one contains more than 98% of the total sequences. ASVs were defined by single-nucleotide differences, with distinct variants identified by a minimum difference of one nucleotide over the sequences (Callahan et al., 2017). The most abundant ASV differs at 16 pair bases compared to *Microdochium seminicola* (NCBI accession number KP859038) but had a 100% match with NCBI accession number MK990150.1 a sequence from maize in Mexico which could be *Microdochium maydis*; however, this comparison is based only on the ITS2 sequences and we have referred to it as *M. seminicola* for simplicity. The isolation of this *Microdochium* from tar spot lesions collected in Ecuador and Guatemala with additional sequence data is required to identify this species conclusively.

*M. seminicola* was detected in both tar spot lesions with fisheye symptoms and those containing only stromata. However, its abundance was higher in lesions with small stromata and fisheye, while it was found in lower abundance in lesions with thick stromata, regardless of the presence of fisheye. For instance, samples from Mocache, in the Coastal Lowlands of Ecuador, which exhibited tar spot lesions with small stromata and fisheye (Fig. 16A), showed a higher abundance of *M. seminicola* compared to *P. maydis*. In contrast, samples from Pueblo Viejo, also in the Coastal Lowlands, where lesions had thick stromata with fisheye (Fig. 16B), showed an abundance of *M. seminicola* of less than 1%.

**Fig. 16.**
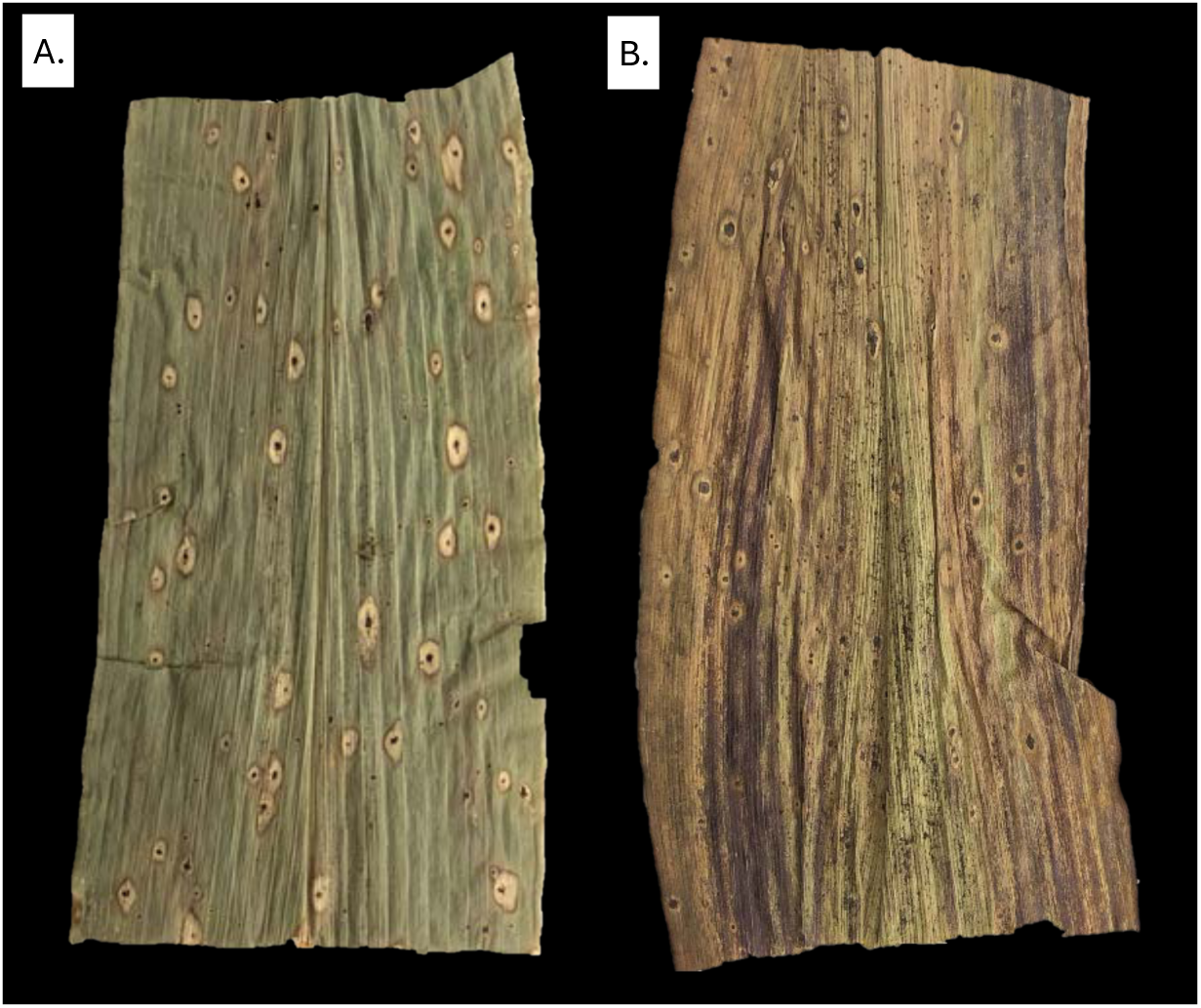
Samples collected from Mocache (A) and Pueblo Viejo (B), located in Coastal Lowlands of Ecuador. Both samples present fisheye symptoms but the abundance of *Microdochium seminicola* in the location Mocache is 68.1% versus 0.06% in Pueblo Viejo.

In Guatemala, fisheye lesions were observed only in Rabinal, Cubulco, and Patzún. Samples from Rabinal and Cubulco exhibited a high abundance of *M. seminicola*, while in Patzún, although two of the three samples had fisheye lesions, the abundance of *M. seminicola* in all samples was less than 2%. Notably, the sample with non-fisheye lesions showed a higher abundance of *M. seminicola* compared to the other two samples. Regardless of the presence of fisheye, the occurrence of *M. seminicola* in tar spot lesions in Ecuador and Guatemala appears to follow a geographical distribution rather than being the cause of fisheye symptoms. Muller and Samuels (1984) reported that *Microdochium maydis* is commonly found in green leaves and considered it an endophyte in corn, with the potential to become pathogenic when in contact with *P. maydis*. Therefore *M. maydis* may be an endophyte that becomes an opportunistic fungus affecting stromata.

*M. seminicola* was found in nearly all collected samples from Guatemala and Ecuador, except for one. The relative abundance of this species was higher than *P. maydis* in some locations in these countries (Table 2). In Ecuador, a high abundance of *M. seminicola* was observed in El Eno and Joya Sachas, located in the Amazon, and Mocache and Vinces from the Coastal Lowlands. It also was high in Cubulco and Rabinal in Guatemala. In all these locations, the relative abundance of *M. seminicola* was higher than *P. maydis* and *Paraphaeosphaeria* sp. In fact, in La Joya Sachas we found *P. maydis* sequences in only two samples from a total of five collected in this location. In this location, samples were collected at the end of the corn season; therefore, all samples were already dry (Fig. 17). The high abundance of *M. seminicola* in tar spot lesions at late infection stages can explain the absence of *P. maydis* sequences due a change of the fungal microbiome over time. Hook et al. (1992) described tar spot lesions after four weeks of infection by microscopy and found that only one-third of the lesions contained ascospores of *P. maydis*. He also found that *Monographella maydis* occurred in 85% of the lesions and *Coniothyrium phyllachorae* in 50% of the lesions. The small size of the stromata also influences the relative abundance of *P. maydis* (Fig. 17). Another possibility is that *Monographella* can cause tar spot-like lesions by itself.

**Fig. 17.**
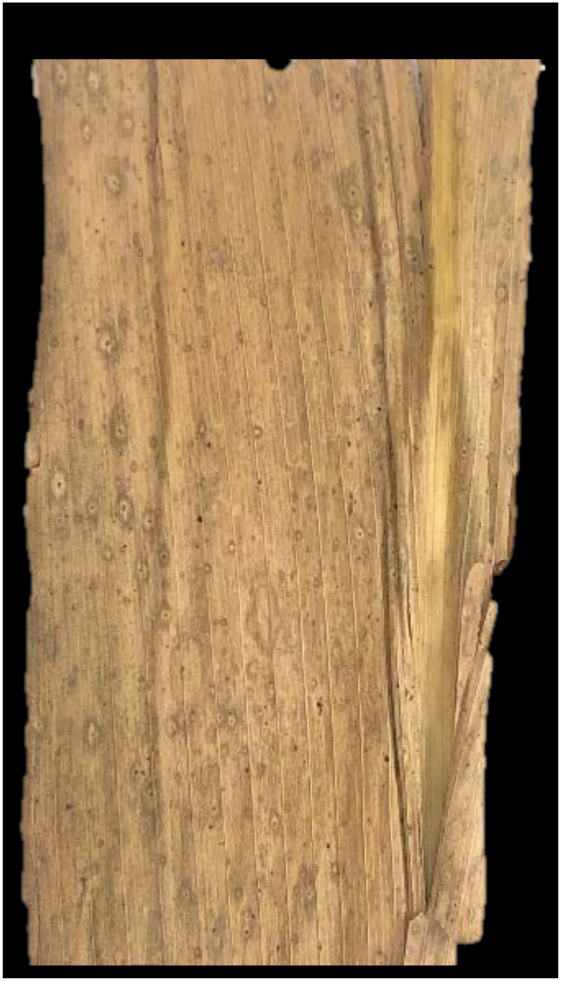
Sample collected from La Joya Sachas, Amazon of Ecuador. Sample was collected when corn leaves were already dry, after corn harvesting. The abundance of *P. maydis* in these samples was low (0.003%) or absent despite the presence of stromata but *Microdochium* was in average 37.7%.

Hock et al. (1992) inoculated the stromata of *P. maydis* with *Microdochium maydis* under laboratory conditions but did not observe infection. They also inoculated leaves with *M. maydis* under field conditions and occasionally observed conidia and sporodochia. They concluded that inoculations with *M. maydis* did not produce infection, but that under unusual conditions, *Microdochium maydis* can infect corn without prior *P. maydis* infections. Therefore, additional experiments are necessary to test the hypothesis that *Monographella maydis* can cause lesions similar to tar spot by itself.

*Alternaria alternata* and *Nigrospora* sp. were the third and fourth most abundant species in USA samples, but their relative abundances were low in Ecuador and Guatemala. *Alternaria alternata* is one of the most common endophytic fungi reported as part of the corn microbiome (Singh & Goodwin, 2022). It also has been reported to be associated with leaf blight of maize in Heilongjiang province of China (Xu et al., 2022). *Nigrospora* is considered a secondary invader, most commonly affecting corn seed and causing kernel rot (Gomaa, 2021). These species appear to be associated mainly with samples that were dried and stored in artisanal presses (see Fig. 18), suggesting that their prevalence may be influenced by storage conditions rather than being integral components of the tar spot microbiome.

**Fig 18.**
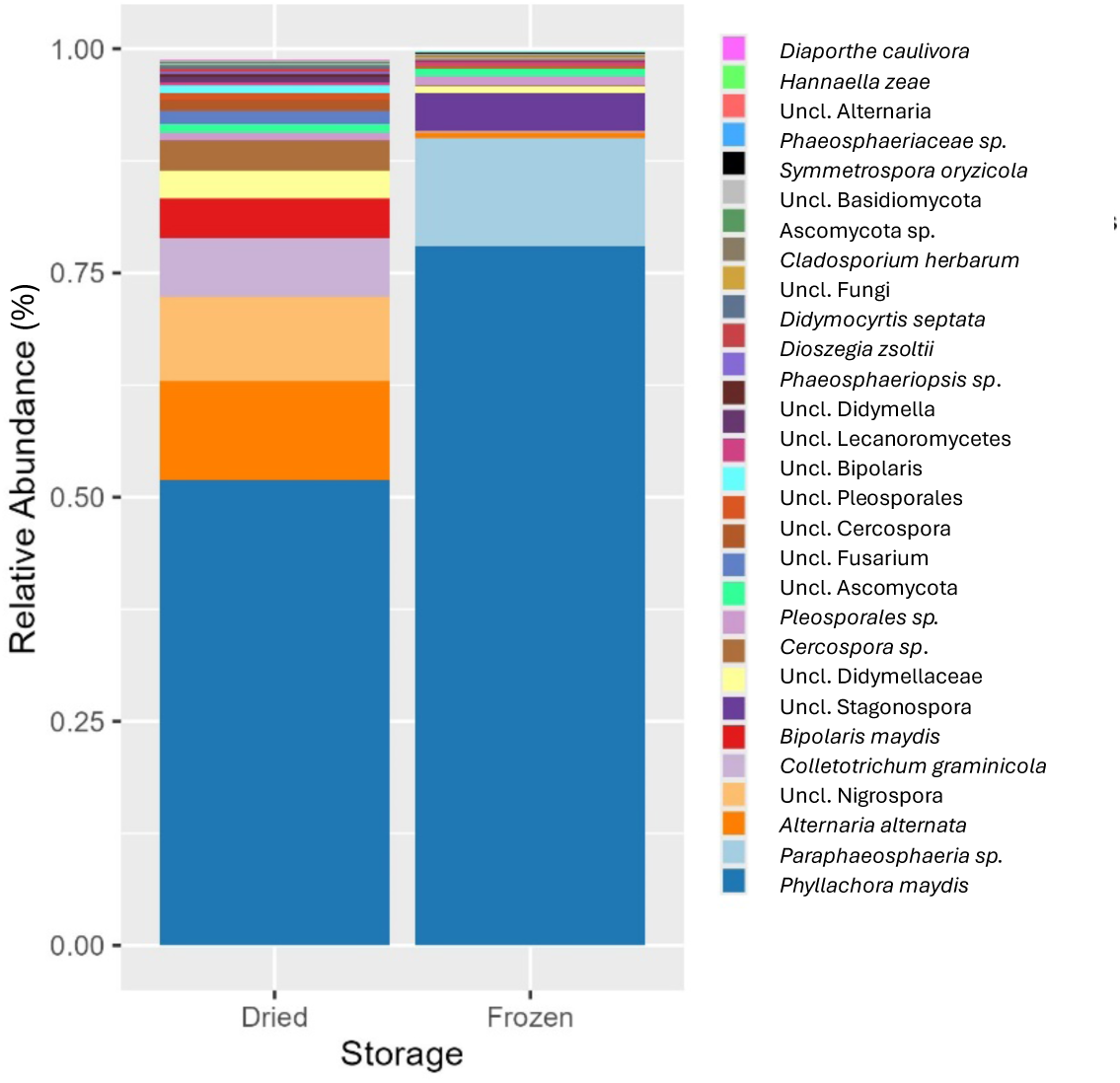
The 30 most abundant fungal species in tar spot lesions collected in Indiana, USA. Samples were collected during the summer of 2021 (dried and stored in a plant press) and in 2023 (frozen at −20°C). DNA extractions were performed in the fall of 2023. Unclassified species belong to a known taxon but could not be identified at the species level. Bars do not reach 100% as species beyond the 30 most abundant are not displayed.

A high relative abundance of reads assigned as *Aspergillus cristatus*, and species within the genus *Aspergillus*, and the family Aspergillaceae, was observed in samples collected in Patzún, Antigua Guatemala, Rabinal, and Cubulco from Guatemala. Our *A. cristatus* sequence (Supplementary Table S7) differs at 11 nucleotide positions compared to the reference sequence in the database, showing 96% similarity (Table 3) so most likely represents a different species. *A. cristatus* is known as the “Golden flower fungus” and is used in the fermentation of *Camellia sinensis* to produce Fu Zhuan tea (Hu et al., 2024); however, we did not find any reports of *A. cristatus* in corn. *Aspergillus* spp. are common fungi in the endospheres of corn but mainly in the seeds (Singh & Goodwin, 2022). Species such as *Aspergillus flavus* produce mycotoxins such as aflatoxins and cause ear rot of corn (Williams et al., 2011). The high abundance of Aspergillus in these locations might be attributed to exposure to some type of plant stress. Maize plants developing under stress conditions, especially during reproductive growth, are predisposed to infection by this fungus and the production of mycotoxins, contaminating the grain (Bruns, 2003). We did not find any relationship between *Aspergillus* spp. and tar spot of corn.

Another taxon in high abundance in some locations in Ecuador is a species placed in the order Russulales. Interestingly, this order is composed of mushroom species. We did not find any previous reports of Russulales on corn and the sequences match with uncultured fungi obtained from environmental samples. Based on unpublished data (M. C. Aime, personal communication), two of the sequences are probably in the same genus of the Peniophoraceae/Lachnocladiaceae as isolates obtained from lichens, leaf surfaces, and cacao pods from Honduras and Guyana, while a third was a 97% match to previously undescribed *Lachnocladium* spp. from Cameroon. Most likely, these Russulales sequences belong to an epiphytic species, but their exact role in the corn microbiome and whether they influence tar spot development remain unknown.

### Fungal microbiome diversity varies across geographical regions and locations

The diversity of the fungal microbiome associated with tar spot of corn exhibits variation across different countries, geographic regions, and specific locations. Samples collected from the USA contained the lowest levels of diversity, whereas diversity in Ecuador and Guatemala varied considerably among locations. This variation may be influenced by several factors, including local environmental conditions, geographic location, climatic factors, corn line, and crop management practices. Samples were collected from geographic regions with different environmental conditions, corn cultivars, and production management practices. All these factors likely shape the fungal microbiome associated with tar spot of corn in each location.

Crop management practices for corn vary significantly among the countries, geographic regions, and specific locations sampled. The production systems in the fields from the USA are more automated and mechanized compared to those in Guatemala and Ecuador, where maize is primarily cultivated under non-commercial systems, particularly in Guatemala. This often involves manual planting and harvesting. Such differences in agricultural practices are known to influence the fungal diversity associated with tar spot in maize. Previous studies have demonstrated that crop management significantly affects the microbiome of sorghum and soybean (Longley et al., 2020; Wipf et al., 2021). For instance, Longley et al. (2020) evaluated the impact of three management systems on the microbiome of soybean across various growth stages, finding that long-term cropping management alters microbial communities in soybean throughout its development. Similarly, Wattenburger et al. (2019) found that agricultural management practices significantly influence the root microbiome of maize, resulting in shifts in community composition as plants develop. These findings highlight the critical role of crop management in shaping microbial diversity in agricultural ecosystems.

Geographic location, local environmental conditions, and host also shape the microbiomes of plants (Christian et al., 2016). This variation in the microbiota is driven by microbial immigration influenced by plant environment. Microbial immigration can occur through aerial deposition, transfer from other plant organs or soil, irrigation, or the application of biological agents (Leveau, 2019). We sampled in total 15 locations in five geographical regions from three countries. Each location has its own environmental conditions and shows a different fungal microbiome composition. Even some locations from the same geographical region showed different fungal microbiome compositions; for example, *Microdochium seminicola* was the predominant species in two locations in the Coastal Lowlands of Ecuador but in the third location from this region this species was almost absent, with *Paraphaeosphaeria* sp. being the most abundant species (Fig. 4).

In Guatemala, the local environment may have influenced the presence and abundance of *Microdochium* and the tar spot microbiome. Samples in this country were collected from the Central Highlands across three states: Chimaltenango, Sacatepéquez, and Baja Verapaz. Chimaltenango and Sacatepéquez are geographically close and share similar environmental conditions. Chimaltenango experiences temperatures ranging from 9.5 to 22.6 °C (49 to 72 °F) with an average relative humidity (RH) of 77%. In contrast, Baja Verapaz exhibits more tropical conditions, with temperatures ranging from 14.4 to 29 °C (57 to 84 °F) and an average RH of 79% (INSIVUMEH, 2022). This variation in environmental conditions affects the corn growth cycle. In temperate regions such as Chimaltenango, the cycle lasts about 8 months, whereas in tropical regions like Baja Verapaz, it lasts only 4 months (Racancoj-Coyoy et al., 2024).

In Guatemala, the corn cycle significantly influences tar spot infection. In Baja Verapaz, tar spot can be detected as early as the first month after planting. If the infection occurs before tasseling in this location, it can result in total production loss. In contrast, in Chimaltenango, where the maize cycle is longer, infection typically occurs after tasseling, about 4–5 months after planting. Consequently, the impact of tar spot is less severe. This highlights the pathogen’s adaptation to the corn cycle in Guatemala (A.J. Racancoj-Coyoy, personal communication).

Other factors that influence fungal microbiomes are crop variety, neighboring vegetation (Whitaker et al., 2023) and leaf age (Geyer et al., 2024). In Indiana, USA, samples were collected from a field planted with a corn hybrid, treated with agrochemicals, and surrounded by large corn fields, while in Guatemala, samples were collected from small plots (less than 1 ha), planted with hybrids, landraces or corn varieties, sometimes surrounded by small plots of corn or forest, with no use of machinery.

The corn growth stage also influences the microbiomes associated with tar spot. The growth stage of the corn plants during sampling ranged from R2 to R6. The three locations in the Coastal Lowlands were sampled at growth stage R2 but, as mentioned earlier, one location differs in fungal composition compared to the other two. The effect of the growth stage on disease development needs to be studied to understand the dynamics of fungal microbiome composition over time in relation to tar spot. This could identify shifts in fungal microbiomes during development and could explain the low abundance of *Phyllachora maydis* in some locations. A shift in fungal microbiomes over time may explain why some samples from the location La Joya Sachas, collected at growth stage R6, did not show any sequences of *P. maydis* despite showing stromata.

Tar spot microbiome samples are separated into four groups, each characterized by different dominant species in the tar spot lesions (Fig. 11). The predominant species in the first group, formed by all samples from Indiana, USA, is *Phyllachora maydis* (Fig. 4). In the second group, which includes all samples from the Ecuadorian Amazon, some locations from the Coastal Lowlands of Ecuador (including Vinces and Mocache), and Guatemala (including Rabinal and Cubulco), the predominant species is *Microdochium seminicola*, while *Phyllachora maydis* and *Paraphaeosphaeria sp.* were found at lower levels. The third group, formed by samples from all locations in the Andean Highlands of Ecuador, some locations from the Coastal Lowlands (including Pueblo Viejo) and Guatemala (including Parramos and Patzún), is dominated by *Phyllachora maydis* and *Paraphaeosphaeria sp.* The last group, comprising samples solely from Guatemala (including Patzún, Técpan, Chimaltenango, and Antigua Guatemala), is dominated by *Phyllachora maydis*, with *Microdochium seminicola* and *Paraphaeosphaeria* sp. at lower levels (Fig. 4). These results suggest that the presence of *Microdochium* and *Paraphaeosphaeria* in tar spot microbiomes follows a geographical distribution but that these species are not associated with tar spot across all locations.

### *Microdochium* is differentially abundant in fisheye samples but is not present in the USA

The high abundance of *Microdochium* in fisheye lesions in Ecuador and Guatemala, along with its differential abundance in fisheye and non-fisheye lesions, supports the hypothesis that this species may play a role in causing fisheye lesions as reported previously (Muller & Samuels, 1984; Sol Hernandez, 2018). However, its absence in fisheye lesions from Indiana, USA, suggests that *Microdochium* is not essential for fisheye lesion development, as suggested by Luis et al. (2023). Instead, our data suggests that its presence may be influenced by geographical factors and corn genotype. In fact, in Guatemala and Ecuador, the abundance of *Microdochium* was low in some locations even when fisheye lesions were observed. Studies have shown that *Microdochium* species do not produce mycotoxins typically associated with filamentous fungi (Gagkaeva et al., 2020; Gavrilova et al., 2020). It appears to be present as an endophyte in non-diseased corn leaves, but Muller & Samuels (1984) suggest that it becomes pathogenic when it comes in contact with *P. maydis*. Controlled inoculations with both species are needed to fully understand their interaction and how it affects tar spot development and fisheye symptoms.

*Phaeosphaeria* and *Curvularia* were also differentially abundant in fisheye lesions. These genera have been proposed as alternative causal agents for fisheye symptoms (McCoy et al., 2019). *Curvularia lunata* has been reported as part of the tar spot complex, along with *P. maydis*, in Mexico (Rios Herrera et al., 2017), and causes leaf spot in corn (Henrickson & Koehler, 2022) and the grassy weed *Hymenachne amplexicaulis* (Monteiro et al., 2003). *Phaeosphaeria maydis* was considered the causal agent of Phaeosphaeria leaf spot (PLS). However, subsequent research identified the bacterium *Pantoea ananatis* as the true cause of PLS (Gonçalves et al., 2013). Interestingly, 76 compounds with diverse biological activity have been isolated from the *Phaeosphaeria* genus (El-Demerdash, 2018), but whether they affect tar spot is not known.

Most of the genera enriched in fisheye lesions exhibit low abundance and are not consistently present across all samples, suggesting limited ecological significance or that they are not causally related to tar spot. Nonetheless, some of these genera can produce secondary metabolites and may possess biological control properties, indicating the need for further research. For example, *Clonostachys*, detected in 24 of the 73 samples, can produce secondary metabolites (Han et al., 2020) and species such as *C. rosea* have been suggested as potential biological controls against fungal plant pathogens, nematodes, and insects (Sun et al., 2020). *Clonostachys* had a relative abundance of 0.41% in fisheye samples and less than 0.01% in tar spot samples, and was primarily found in Ecuador and Guatemala (Fig 5). *Sarocladium* species, found in 25 of the 73 samples, are usually associated with grasses as saprobes, parasites, or endophytes (Yeh & Kirschner, 2014). For example, *Sarocladium oryzae* can cause sheath rot disease in rice (Pramunadipta et al., 2020) and *S. zeae* acts as an endophyte in wheat (Kemp et al., 2020). Both of these species can produce secondary metabolites (Błaszczyk et al., 2021), phytotoxic metabolites (Sakthivel et al., 2002), or act as biocontrol agents (Kemp et al., 2020).

The basidiomycetous yeast *Hannaella* was found in 71 of the 73 samples, with a relative abundance of 0.9% in fisheye samples and 0.7% in tar spot samples (Fig 5). This taxon, which includes species from the *Bullera sinensis* clade, is widely distributed on plant leaf surfaces and plays a significant role as a phyllosphere-inhabiting yeast (Landell et al., 2014; Q.-M. Wang & Bai, 2008). *Hannaella sinensis* has been reported in corn plants in Thailand (Into et al., 2020) and has shown potential as a biocontrol agent, inhibiting the growth of *Aspergillus flavus* mycelia through the production of volatile organic compounds (VOCs) (Jaibangyang et al., 2020). Neto et al. (2021) showed that yeasts living in aerial parts of *Theobroma cacao* plants are prospective biological control agents due to their ability to produce killer toxins, volatile compounds, and hydrolytic enzymes that can inhibit the growth of other organisms. However, they also showed that not all *Hannaella* spp. exhibit the desired traits for a BCA agent.

Interestingly, some of the genera enriched in lesions with non-fisheye symptoms are also known to produce leaf spots. For example, species in the family Mycosphaerellaceae, a taxon found in 28 out of 73 samples, can cause leaf spots in Proteaceae species (Taylor et al., 2003), while a *Neoascochyta* species, found in 26 out of 73 samples, has been identified as a cause of leaf scorch on wheat (Golzar & Wang, 2012). Additionally, the species *Phaeosphaeriopsis glaucopunctata*, a genus found in 56 out of 73 samples, causes leaf spot and necrosis on Butcher’s Broom (*Ruscus aculeatus*) (Golzar & Wang, 2012). The genus *Neosetophoma*, found in 42 out of 73 samples, was created to accommodate the *Phoma samarorum* clade that causes leaf spot on grasses (De Gruyter et al., 2010; Marin-Felix et al., 2019). Notably, *N. poaceicola* was isolated from black spot or scab-like symptoms in apples (Ebrahimi & Fotouhifar, 2021). The differential abundance of these genera in fisheye versus non-fisheye lesions is also low and they were not present in all samples.

The abundance of *Alternaria* was higher compared to other differentially abundant genera. *Alternaria* is present in all samples except one from Guatemala (Fig 5), with a relative abundance of 0.58% in fisheye samples and 4.5% in non-fisheye tar spot samples. A*lternaria* is commonly found in corn and acts as an opportunistic pathogen, infecting the host after damage caused by other fungi. Furthermore, it has been found in high abundance in both asymptomatic tissue and typical tar spot with non-fisheye symptoms (relative abundance of 20.7% and 19.0%, respectively) but occurred in low abundance in fisheye lesions (relative abundance of 6.1%) (McCoy et al., 2019). Additionally, *Alternaria* species recovered from overwintered tar spot stromata were tested for their ability to reduce the number of stromata that could produce ascospores and were proposed as a potential biological control (Johnson et al., 2023). Another genus with antagonistc proprieties is *Bullera alba*, which has been reported as an endophytic antagonist yeast against gray mold (*Botrytis cinerea*) in apple (Yu et al., 2024). *Bullera* was found in 11 of 40 fisheye samples and 17 of 33 non-fisheye tar spot samples, mostly from the USA, with a relative abundance of less than 0.1% in both types of samples.

The differential analysis of the microbiomes identified 14 genera that were enriched in fisheye relative to non-fisheye lesions. However, most of these genera are characterized by their capacity to produce secondary metabolites, suggesting possible use as biological control agents, rather than as species that cause necrotic lesions. The role of *Microdochium* in tar spot is still unclear, whether it causes fisheye lesions, exists as an endophyte within corn leaves, or can cause tar spot-like lesions by itself in Ecuador and Guatemala.

### Correlations between *P. maydis* and other fungi

Most correlations involving *P. maydis* were negative, except for the likely mycoparasite *Paraphaeosphaeria* sp. This species has been identified within tar spot stromata (Caldwell et al., 2023) and several species that previously were considered *Coniothyrium* have been reclassified under *Paraphaeosphaeria* (Verkley et al., 2014). A positive correlation could occur if this species mostly functions as a mycoparasite, with its frequency increasing in proportion to its food source. Thus, further investigation into *Paraphaeosphaeria* sp. is warranted to assess its potential as a biological control agent against *P. maydis*. Interestingly, the correlation between *P. maydis* and *Coniothyrium sidae*, another possible mycoparasite, is negative; however, the abundance of *C. sidae* was low, less than 0.02% per sample, and inconsistent, being present in only 16 out of 73 samples.

Other ASVs positively correlated with *P. maydis* included some that could not be classified at the species level. These ASVs were identified as members of the families Didymellaceae and Sarocladiaceae, the class Dothideomycetes, and the genera *Cercospora* and *Fusarium*. These taxa comprise a wide range of species, genera, families, and classes with substantial diversity, making it challenging to generalize their roles in relation *to P. maydis*. Furthermore, the BLAST results (Table 3) were inconclusive, as the sequences matched multiple species within the same genus, all with identical scores, e-values, and identity percentages. For example, the most abundant ASVs classified within Didymellaceae matched species from both genera *Epicoccum* and *Allophoma*.

The frequency of *P. maydis* was negatively correlated with *Phaeosphaeria oryzae*, *Microdochium seminicola,* and a species in the genus *Nigrospora*. *Nigrospora* is a cosmopolitan genus that has been attributed with roles as an endophyte, saprobe, plant pathogen, or opportunistic human pathogen (Wang et al., 2017). Species such as *N. sphaerica*, which is known to cause plant diseases, can produce compounds with fungicidal properties that are able to inhibit the mycelial growth of the Oomycete *Phytophthora infestans* (Kim et al., 2001). *Phaeosphaeria* can produce phaeofungin, a compound that has been shown to have antifungal activity against *Candida albicans*, *Aspergillus fumigatus* and *Trichophyton mentagrophytes* (Singh et al., 2013). Therefore, the negative correlation of this species with *P. maydis* could be a result of the production of this antifungal compound. Interestingly, *Phaeosphaeria oryzae* exhibits a negative correlation with *Paraphaeosphaeria* sp., which could indicate competition for food resources if both species act as mycoparasites. Therefore, these species also need to be further studied to understand their role in tar spot of corn and their potential as biological control agents.

The negative correlation between *P. maydis* and *Microdochium*, in addition to the observed high abundance of *Microdochium* and low or no presence of *P. maydis* in some samples, suggests that *Microdochium* can produce tar spot-like lesions in some locations of Ecuador and Guatemala. If this is correct, it may represent a new disease with *Microdochium* as a primary pathogen. Isolation of this fungus in culture and extensive inoculation experiments are needed to test this hypothesis.

## CONCLUSIONS

The frequencies of *Phyllachora maydis*, *Paraphaeosphaeria* sp., and *Microdochium seminicola* are highly variable among tar spot lesions in the USA, Ecuador, and Guatemala. Their abundance and presence seem to be driven by geographical location, and there is no single species that is always present within tar spot lesions. The stroma size and presence of fisheye symptoms are associated with the presence of *Paraphaeosphaeria* and *Microdochium*. The tar spot microbiome can be categorized into four distinct groups, each characterized by a species that predominates in tar spot lesions. In the first two groups, one containing samples from Guatemala and the other from Indiana, USA, *Phyllachora maydis* is the most abundant species. A third cluster is dominated by both *P. maydis* and *Paraphaeosphaeria*, comprising samples from Educator and Guatemala. The fourth cluster, which includes samples from Ecuador and Guatemala, is characterized by the predominance of *Microdochium* in higher abundance than *P. maydis* with some samples almost lacking the latter species entirely. This study confirms the presence of *Microdochium* in tar spot lesions outside the USA and suggests that its association with tar spot is dependent on geographical location and that it could cause tar spot-like lesions in Ecuador and Guatemala rather than being a causative factor for fisheye symptoms.

## ACKNOWLEDGMENTS

Sampling in Indiana was possible to Purdue University as part of AgSEED Crossroads funding to support Indiana’s Agriculture and Rural Development.

## Funding

This work was supported by USDA-ARS research project 401-5020-050 014-000D, and Purdue startup funds allocated to Dr. C. D. Cruz.

## Supplementary materials

**Supplementary Fig S1.**
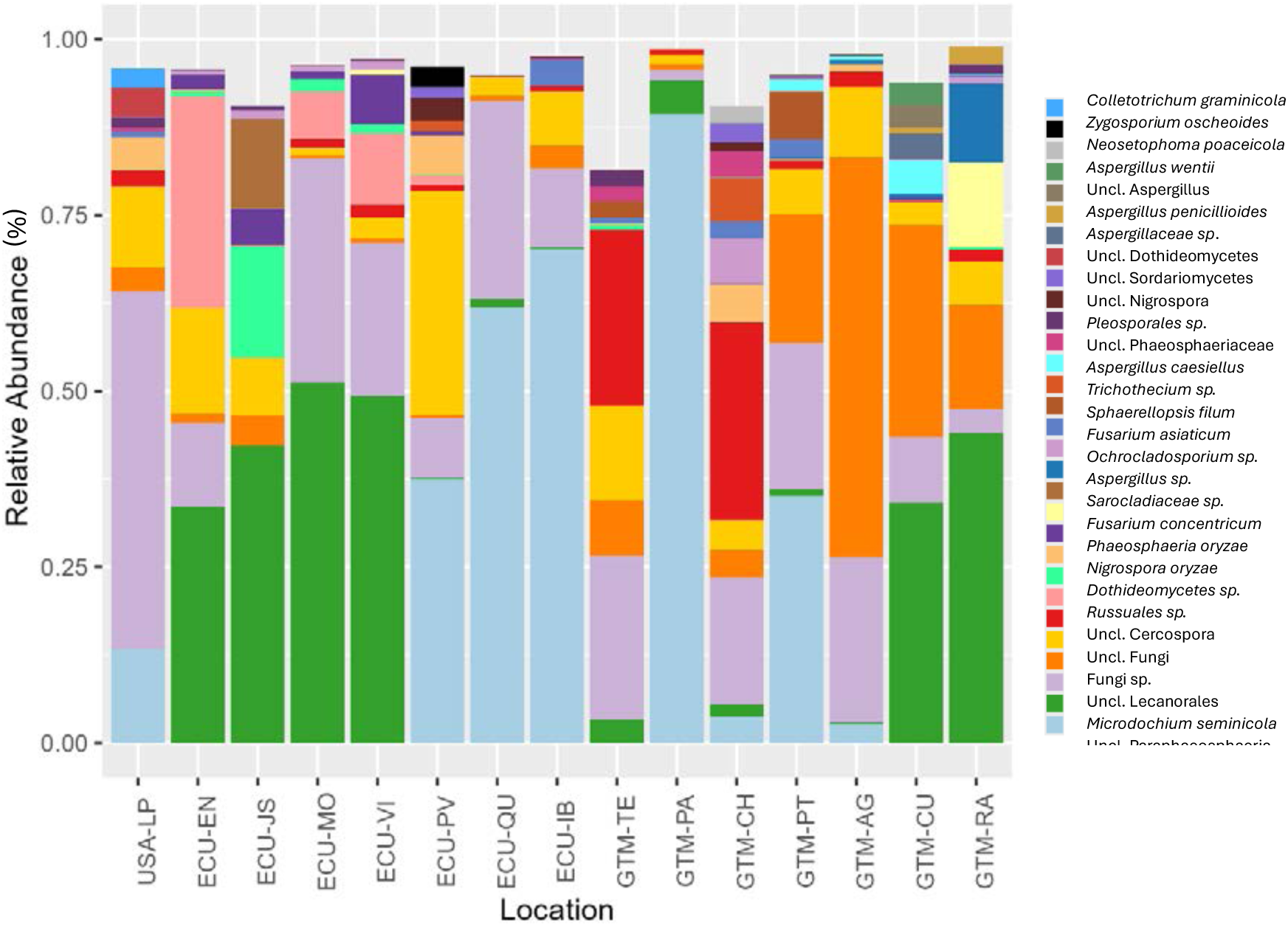
The 30 most abundant fungal species in tar spot lesions by location using ITS1 sequencing reads. ITS1 sequences did not provide enough taxonomic resolution to identify *P. maydis*. USA-LP: La Porte, Indiana; ECU-EN: El Eno, Ecuadorian Amazon; ECU-JS: La Joya Sachas, Ecuadorian Amazon; ECU-MO: Mocache, Coastal Lowlands of Ecuador; ECU-VI: Vinces, Coastal Lowlands of Ecuador; ECU-PV: Pueblo Viejo, Coastal Lowlands of Ecuador; ECU-QU: Quito, Andean Higlands of Ecuador; ECU-IB: Ibarra, Andean Highlands of Ecuador; GTM-TE: Técpan, Guatemala. GTM-PA: Parramos, Guatemala; GTM-CH: Chimaltenango, Guatemala; GTM-PT: Patzún, Guatemala; GTM-AG: Antigua Guatemala, Guatemala; GTM-CU: Cubulco, Guatemala and GTM-RA: Rabinal, Guatemala. Uncl.: Unclassified species that belong to a taxon but could not be assigned to a specific species. Bars do not reach 100% as species beyond the 30 most abundant are not displayed.

**Supplementary Fig S2.**
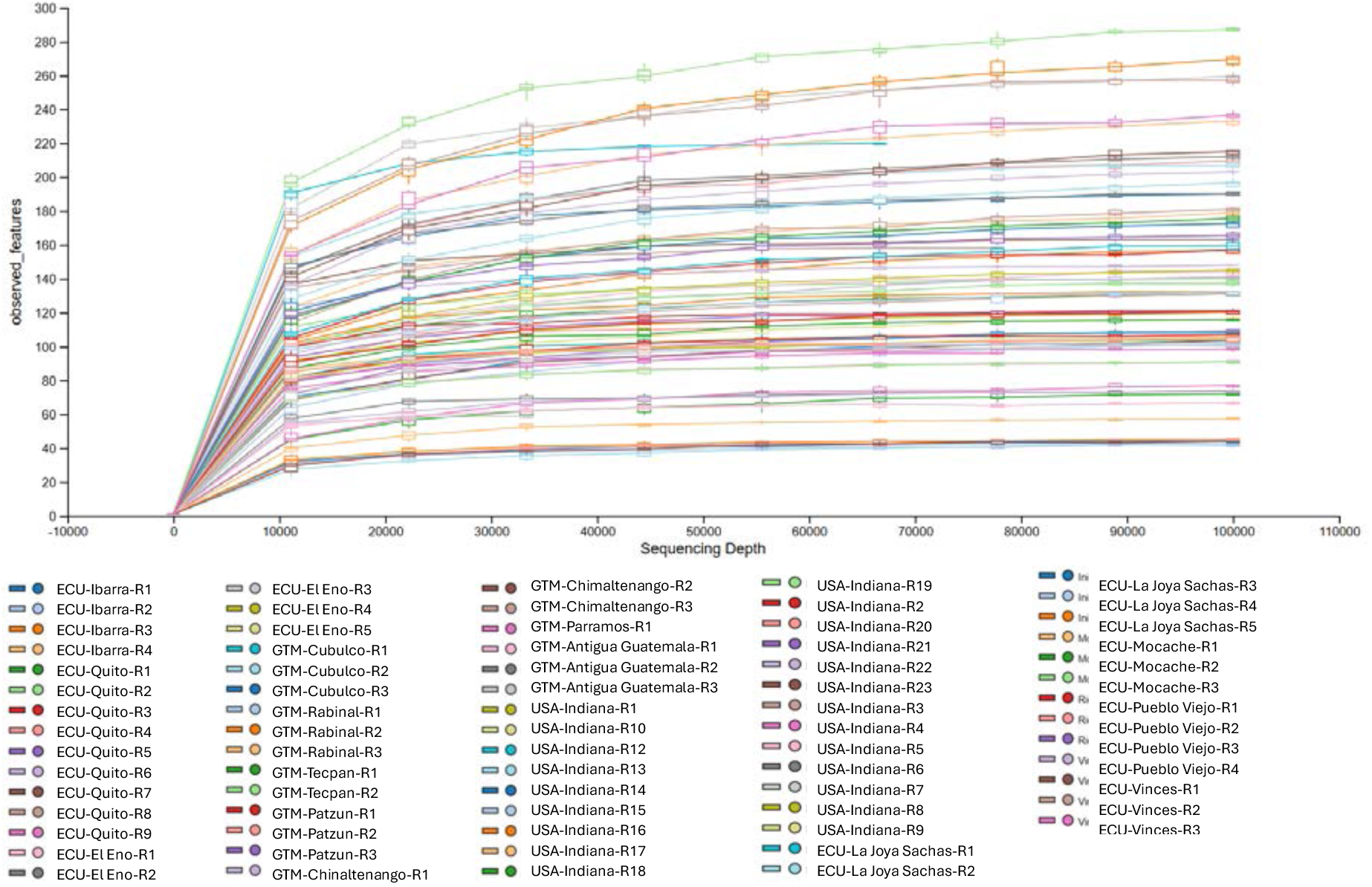
Rarefaction curves of tar spot microbiome samples collected in different locations from Ecuador (ECU), Guatemala (GTM) and USA (United States). The sample GTM-Antigua Guatemala-3 has 41,418 reads, which is the lowest number of reads per sample. At this sequencing depth, the other samples began to reach a plateau.

**Supplementary Table S1.** Global tests for alpha diversity indices compare diversity across groups categorized by countries and regions. The Kruskal-Wallis rank sum test evaluates whether the medians of the diversity indices differ significantly among the groups.

| Index | Comparison Type | Groups | Chi-Squared | Degrees of Freedom | p-value |
| --- | --- | --- | --- | --- | --- |
| Pielou's Evenness | Countries | Guatemala, Ecuador, USA | 11.461 | 2 | 0.0032 |
|  | Regions | USA: Indiana; Guatemala: Central Highlands; Ecuador: Amazon, Coastal Lowlands, Andean Highlands | 25.295 | 4 | 0.0000439 |
| Faith Phylogenetic | Countries | Guatemala, Ecuador, USA | 28.595 | 2 | 0.000000618 |
|  | Regions | USA: Indiana; Guatemala: Central Highlands; Ecuador: Amazon, Coastal Lowlands, Andean Highlands | 32.056 | 4 | 0.00000186 |
| Shannon | Countries | Guatemala, Ecuador, USA | 14.844 | 2 | 0.0005981 |
|  | Regions | USA: Indiana; Guatemala: Central Highlands; Ecuador: Amazon, Coastal Lowlands, Andean Highlands | 29.728 | 4 | 0.00000556 |

**Supplementary Table S2.** Pairwise comparisons of alpha diversity indices between countries using Wilcoxon rank-sum tests with Benjamini-Hochberg (BH) p-value adjustment. Statistically significant p-values (e.g., < 0.05) indicate differences in diversity between the groups compared.

| Index | Group 1 | Group 2 | p-value |
| --- | --- | --- | --- |
| Pielou's Evenness | Guatemala | USA | 0.004 |
|  |  | Ecuador | 0.052 |
|  | Ecuador | USA | 0.5 |
| Faith Phylogenetic | Guatemala | USA | 0.02954 |
|  |  | Ecuador | 0.00017 |
|  | Ecuador | USA | 0.0000012 |
| Shannon | Guatemala | USA | 0.0013 |
|  |  | Ecuador | 0.1326 |
|  | Ecuador | USA | 0.0032 |

**Supplementary Table S3.** Pairwise comparisons of alpha diversity indices between regions using Wilcoxon rank-sum tests with Benjamini-Hochberg (BH) p-value adjustment. Statistically significant p-values (e.g., < 0.05) indicate differences in diversity between the groups compared.

| Index | Group 1 | Group 2 | p-value |
| --- | --- | --- | --- |
| Pielou's Evenness | Central Highlands<br>Guatemala | Indiana - USA | 0.0045 |
|  |  | Amazon - Ecuador | 0.6611 |
|  |  | Coastal Lowlands - Ecuador | 0.7237 |
|  |  | Andean Highlands - Ecuador | 0.000013 |
|  | Andean Highlands<br>Ecuador | Indiana - USA | 0.7237 |
|  |  | Amazon - Ecuador | 0.000013 |
|  |  | Coastal Lowlands - Ecuador | 0.0049 |
|  | Coastal Lowlands<br>Ecuador | Indiana-USA | 0.0713 |
|  |  | Amazon -Ecuador | 0.6611 |
|  | Amazon<br>Ecuador | Indiana - USA | 0.0186 |
| Faith Phylogenetic | Central Highlands<br>Guatemala | Indiana - USA | 0.0422 |
|  |  | Amazon - Ecuador | 0.00025 |
|  |  | Coastal Lowlands – Ecuador | 0.0028 |
|  |  | Andean Highlands – Ecuador | 0.06384 |
|  | Andean Highlands<br>Ecuador | Indiana-USA – Ecuador | 0.0028 |
|  |  | Amazon – Ecuador | 0.0422 |
|  |  | Coastal Lowlands – Ecuador | 0.06384 |
|  | Coastal Lowlands<br>Ecuador | Indiana - USA | 0.00025 |
|  |  | Amazon – Ecuador | 0.9118 |
|  | Amazon<br>Ecuador | Indiana - USA | 0.00025 |
| Shannon | Central highlands<br>Guatemala | Indiana - USA | 0.0011 |
|  |  | Amazon – Ecuador | 0.4407 |
|  |  | Coastal Lowlands – Ecuador | 0.9812 |
|  |  | Andean Highlands – Ecuador | 0.000099 |
|  | Andean Highlands<br>Ecuador | Indiana - USA | 0.5102 |
|  |  | Amazon – Ecuador | 0.000017 |
|  |  | Coastal Lowlands – Ecuador | 0.003 |
|  | Coastal Lowlands<br>Ecuador | Indiana - USA | 0.0067 |
|  |  | Amazon – Ecuador | 0.7006 |
|  | Amazon<br>Ecuador | Indiana - USA | 0.0011 |

**Supplementary Table S4.** PERMANOVA results of weighted UniFrac and Bray Curtis diversity indices based on country and region comparisons. R² indicates the proportion of variance explained, and p-values were adjusted using the BH method.NA indicates empty cells.

| Index | Df | Sum of Sqs | R<sup>2</sup> | F | Pr(>F) |
| --- | --- | --- | --- | --- | --- |
| Bray Curtis dissimilarity |  |  |  |  |  |
| Country | 2 | 9.51 | 0.33 | 17.52855 | 0.001 |
| Residual | 70 | 18.99 | 0.67 | NA | NA |
| Total | 72 | 28.51 | 1 | NA | NA |
| Region | 4 | 14.26 | 0.50 | 17.01938 | 0.001 |
| Residual | 68 | 14.24 | 0.50 | NA | NA |
| Total | 72 | 28.51 | 1 | NA | NA |
| weighted UniFrac distance |  |  |  |  |  |
| Country | 2 | 3.89 | 0.24 | 10.85 | 0.001 |
| Residual | 70 | 12.56 | 0.76 | NA | NA |
| Total | 72 | 16.46 | 1 | NA | NA |
| Region | 4 | 6.65 | 0.40 | 11.53 | 0.001 |
| Residual | 68 | 9.80 | 0.60 | NA | NA |
| Total | 72 | 16.46 | 1 | NA | NA |

**Supplementary Table S5.**
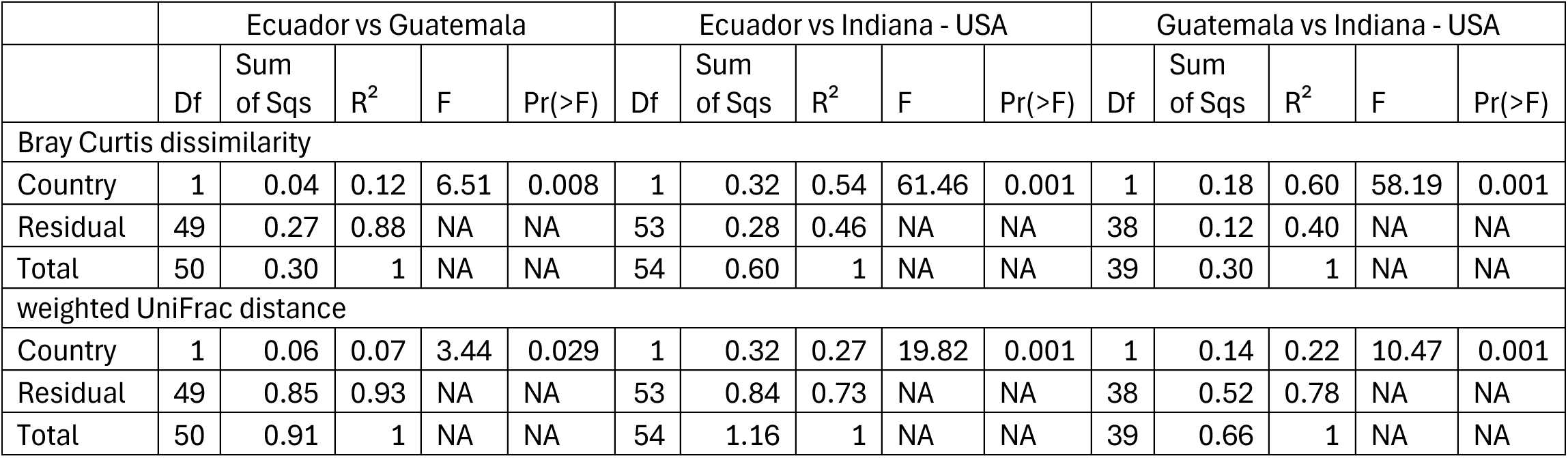
Statistical results of pairwise permutation tests comparing different countries. Comparisons were performed using 999 permutations. NA indicates empty cells.

|  | Ecuador vs Guatemala |  |  |  |  | Ecuador vs Indiana - USA |  |  |  |  | Guatemala vs Indiana - USA |  |  |  |  |
| --- | --- | --- | --- | --- | --- | --- | --- | --- | --- | --- | --- | --- | --- | --- | --- |
|  | Df | Sum of Sqs | R<sup>2</sup> | F | Pr(>F) | Df | Sum of Sqs | R<sup>2</sup> | F | Pr(>F) | Df | Sum of Sqs | R<sup>2</sup> | F | Pr(>F) |
| Bray Curtis dissimilarity |  |  |  |  |  |  |  |  |  |  |  |  |  |  |  |
| Country | 1 | 0.04 | 0.12 | 6.51 | 0.008 | 1 | 0.32 | 0.54 | 61.46 | 0.001 | 1 | 0.18 | 0.60 | 58.19 | 0.001 |
| Residual | 49 | 0.27 | 0.88 | NA | NA | 53 | 0.28 | 0.46 | NA | NA | 38 | 0.12 | 0.40 | NA | NA |
| Total | 50 | 0.30 | 1 | NA | NA | 54 | 0.60 | 1 | NA | NA | 39 | 0.30 | 1 | NA | NA |
| weighted UniFrac distance |  |  |  |  |  |  |  |  |  |  |  |  |  |  |  |
| Country | 1 | 0.06 | 0.07 | 3.44 | 0.029 | 1 | 0.32 | 0.27 | 19.82 | 0.001 | 1 | 0.14 | 0.22 | 10.47 | 0.001 |
| Residual | 49 | 0.85 | 0.93 | NA | NA | 53 | 0.84 | 0.73 | NA | NA | 38 | 0.52 | 0.78 | NA | NA |
| Total | 50 | 0.91 | 1 | NA | NA | 54 | 1.16 | 1 | NA | NA | 39 | 0.66 | 1 | NA | NA |

**Supplementary Table S6.**
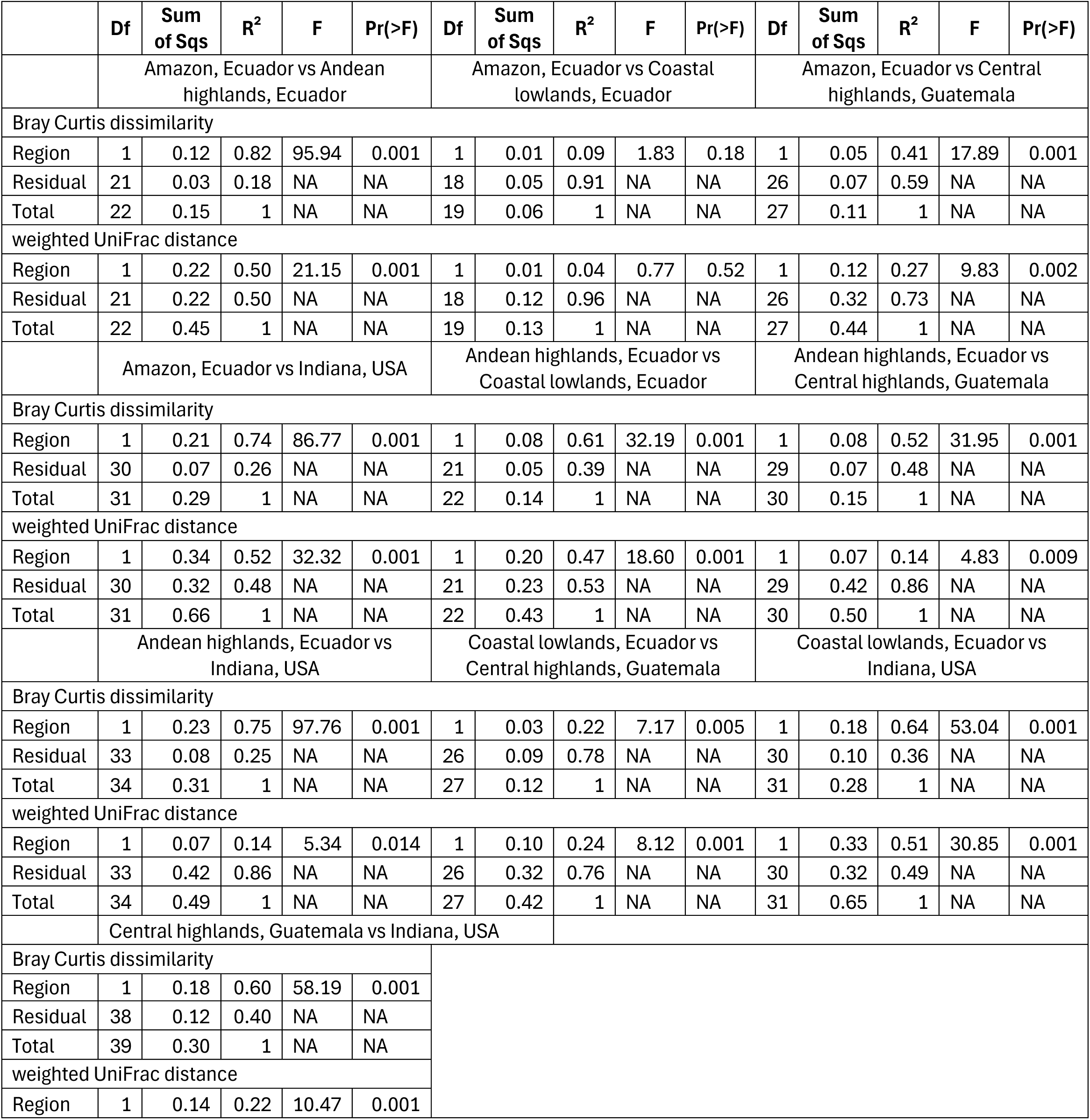
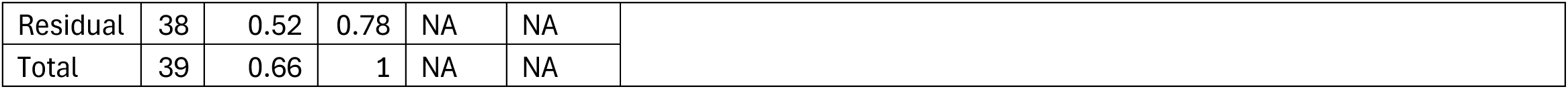
Statistical results of pairwise permutation tests comparing different regions. Comparisons were performed using 999 permutations. NA indicates empty cells.

|  | Df | Sum of Sqs | R<sup>2</sup> | F | Pr(>F) | Df | Sum of Sqs | R<sup>2</sup> | F | Pr(>F) | Df | Sum of Sqs | R<sup>2</sup> | F | Pr(>F) |
| --- | --- | --- | --- | --- | --- | --- | --- | --- | --- | --- | --- | --- | --- | --- | --- |
|  | Amazon, Ecuador vs Andean highlands, Ecuador |  |  |  |  | Amazon, Ecuador vs Coastal lowlands, Ecuador |  |  |  |  | Amazon, Ecuador vs Central highlands, Guatemala |  |  |  |  |
| Bray Curtis dissimilarity |  |  |  |  |  |  |  |  |  |  |  |  |  |  |  |
| Region | 1 | 0.12 | 0.82 | 95.94 | 0.001 | 1 | 0.01 | 0.09 | 1.83 | 0.18 | 1 | 0.05 | 0.41 | 17.89 | 0.001 |
| Residual | 21 | 0.03 | 0.18 | NA | NA | 18 | 0.05 | 0.91 | NA | NA | 26 | 0.07 | 0.59 | NA | NA |
| Total | 22 | 0.15 | 1 | NA | NA | 19 | 0.06 | 1 | NA | NA | 27 | 0.11 | 1 | NA | NA |
| weighted UniFrac distance |  |  |  |  |  |  |  |  |  |  |  |  |  |  |  |
| Region | 1 | 0.22 | 0.50 | 21.15 | 0.001 | 1 | 0.01 | 0.04 | 0.77 | 0.52 | 1 | 0.12 | 0.27 | 9.83 | 0.002 |
| Residual | 21 | 0.22 | 0.50 | NA | NA | 18 | 0.12 | 0.96 | NA | NA | 26 | 0.32 | 0.73 | NA | NA |
| Total | 22 | 0.45 | 1 | NA | NA | 19 | 0.13 | 1 | NA | NA | 27 | 0.44 | 1 | NA | NA |
|  | Amazon, Ecuador vs Indiana, USA |  |  |  |  | Andean highlands, Ecuador vs Coastal lowlands, Ecuador |  |  |  |  | Andean highlands, Ecuador vs Central highlands, Guatemala |  |  |  |  |
| Bray Curtis dissimilarity |  |  |  |  |  |  |  |  |  |  |  |  |  |  |  |
| Region | 1 | 0.21 | 0.74 | 86.77 | 0.001 | 1 | 0.08 | 0.61 | 32.19 | 0.001 | 1 | 0.08 | 0.52 | 31.95 | 0.001 |
| Residual | 30 | 0.07 | 0.26 | NA | NA | 21 | 0.05 | 0.39 | NA | NA | 29 | 0.07 | 0.48 | NA | NA |
| Total | 31 | 0.29 | 1 | NA | NA | 22 | 0.14 | 1 | NA | NA | 30 | 0.15 | 1 | NA | NA |
| weighted UniFrac distance |  |  |  |  |  |  |  |  |  |  |  |  |  |  |  |
| Region | 1 | 0.34 | 0.52 | 32.32 | 0.001 | 1 | 0.20 | 0.47 | 18.60 | 0.001 | 1 | 0.07 | 0.14 | 4.83 | 0.009 |
| Residual | 30 | 0.32 | 0.48 | NA | NA | 21 | 0.23 | 0.53 | NA | NA | 29 | 0.42 | 0.86 | NA | NA |
| Total | 31 | 0.66 | 1 | NA | NA | 22 | 0.43 | 1 | NA | NA | 30 | 0.50 | 1 | NA | NA |
|  | Andean highlands, Ecuador vs Indiana, USA |  |  |  |  | Coastal lowlands, Ecuador vs Central highlands, Guatemala |  |  |  |  | Coastal lowlands, Ecuador vs Indiana, USA |  |  |  |  |
| Bray Curtis dissimilarity |  |  |  |  |  |  |  |  |  |  |  |  |  |  |  |
| Region | 1 | 0.23 | 0.75 | 97.76 | 0.001 | 1 | 0.03 | 0.22 | 7.17 | 0.005 | 1 | 0.18 | 0.64 | 53.04 | 0.001 |
| Residual | 33 | 0.08 | 0.25 | NA | NA | 26 | 0.09 | 0.78 | NA | NA | 30 | 0.10 | 0.36 | NA | NA |
| Total | 34 | 0.31 | 1 | NA | NA | 27 | 0.12 | 1 | NA | NA | 31 | 0.28 | 1 | NA | NA |
| weighted UniFrac distance |  |  |  |  |  |  |  |  |  |  |  |  |  |  |  |
| Region | 1 | 0.07 | 0.14 | 5.34 | 0.014 | 1 | 0.10 | 0.24 | 8.12 | 0.001 | 1 | 0.33 | 0.51 | 30.85 | 0.001 |
| Residual | 33 | 0.42 | 0.86 | NA | NA | 26 | 0.32 | 0.76 | NA | NA | 30 | 0.32 | 0.49 | NA | NA |
| Total | 34 | 0.49 | 1 | NA | NA | 27 | 0.42 | 1 | NA | NA | 31 | 0.65 | 1 | NA | NA |
|  | Central highlands, Guatemala vs Indiana, USA |  |  |  |  |  |  |  |  |  |  |  |  |  |  |
| Bray Curtis dissimilarity |  |  |  |  |  |  |  |  |  |  |  |  |  |  |  |
| Region | 1 | 0.18 | 0.60 | 58.19 | 0.001 |  |  |  |  |  |  |  |  |  |  |
| Residual | 38 | 0.12 | 0.40 | NA | NA |  |  |  |  |  |  |  |  |  |  |
| Total | 39 | 0.30 | 1 | NA | NA |  |  |  |  |  |  |  |  |  |  |
| weighted UniFrac distance |  |  |  |  |  |  |  |  |  |  |  |  |  |  |  |
| Region | 1 | 0.14 | 0.22 | 10.47 | 0.001 |  |  |  |  |  |  |  |  |  |  |

|  |  |  |  |  |  |
| --- | --- | --- | --- | --- | --- |
| Residual | 38 | 0.52 | 0.78 | NA | NA |
| Total | 39 | 0.66 | 1 | NA | NA |

## Notes

### Competing Interest Statement

The authors have declared no competing interest.

